# CD45 sequestration lowers the signaling threshold in lymphocytes and enhances anti-tumor immunity

**DOI:** 10.1101/2025.07.29.667400

**Authors:** Lauren Duhamel, Yiming J. Zhang, William Pinney, Elizabeth Fink, Qingyang Henry Zhao, Anna Romanov, Jordan A. Stinson, Luciano Santollani, Joseph R. Palmeri, Owen T. Porth, Darrell J. Irvine, K. Dane Wittrup

**Affiliations:** Department of Biological Engineering, Massachusetts Institute of Technology, Cambridge, MA, USA; Koch Institute for Integrative Cancer Research, Massachusetts Institute of Technology, Cambridge, MA, USA; Department of Chemical Engineering, Massachusetts Institute of Technology, Cambridge, MA, USA; Howard Hughes Medical Institute, Chevy Chase, MD, USA; Department of Immunology & Microbiology, The Scripps Research Institute, San Diego, CA, USA

## Abstract

CD45 plays a central role in immune signal regulation by controlling the spatial dynamics of phosphatase activity through steric segregation of its bulky rigid extracellular domain. To modulate CD45 activity, here we develop and characterize protein engineering approaches to induce multivalent clustering of CD45, effectively mimicking the endogenous local receptor sequestration during immune synapse formation. In doing so, we engineer a biologic that enables precise, tunable control over CD45 surface localization and activity. CD45 sequestration exhibited striking synergy when administered in combination with intratumorally anchored IL-12 therapy, markedly delaying tumor progression and extending survival in syngeneic murine melanoma and carcinoma models. Immune profiling revealed that CD8⁺ T cells are essential mediators of this synergistic antitumor response. Mechanistically, IL-12 initiates a wave of antigen generation and T cell priming, while CD45 sequestration subsequently enhances tumor-specific CD8⁺ T cell activation, expansion, and functional states within the tumor-draining lymph node. These findings suggest that CD45 sequestration lowers the activation threshold of T cells, broadens the tumor-reactive T cell repertoire, and therefore promotes more robust tumor-specific T cell responses. Altogether, we establish CD45 as a promising novel target for cancer immunotherapy, capable of potentiating strong anticancer immune responses.

## Main Text

CD45 is a receptor protein tyrosine phosphatase highly expressed on the surface of all leukocytes, and has long been implicated as an essential regulator of key signal transduction pathways in immune cells^1^. Its phosphatase activity is regulated via steric segregation of its large, rigid extracellular domain, as described by the kinetic segregation model^2,3^. Immune receptor signaling is amplified when CD45 is physically excluded from sites of receptor engagement, allowing localized phosphorylation by Src-family kinases to proceed unopposed. Signal regulation by CD45 kinetic exclusion has been documented across a wide range of lymphocyte subtypes and receptor families including the T cell receptor (TCR)^4–7^, killer cell immunoglobulin-like receptors^8–10^, phagocytic receptors^11,12^, and the immunoglobulin E receptor^13^. Beyond native receptor-ligand interactions, CD45 spatial segregation has also been shown to influence the activity of chimeric antigen receptor (CAR) T cells and agonist antibodies^14,15^.

The mechanism of CD45 regulation by steric segregation is most well characterized in the context of TCR - peptide major histocompatibility complex (pMHC) interactions^4–7^. TCR signaling is governed by a finely tuned balance of kinases and phosphatases, with CD45 playing a key role by modulating the activity of the Src-family kinase Lck. CD45 maintains the basal activity of Lck by dephosphorylating the inhibitory Y505 residue. During T cell activation and TCR triggering, exclusion of CD45 from sites of TCR–pMHC engagement prevents dephosphorylation of the activating Y394 tyrosine on Lck, allowing accumulation of active Lck to initiate downstream signaling. Under conditions of weak antigen stimulation, CD45 acts as a dominant suppressor of low-avidity T cell responses^16,17^. Strategies to transiently attenuate CD45 activity therefore represent a promising avenue to enhance T cell activation in settings where immune responses are otherwise suboptimal, for example in the context of cancer or chronic infection. Despite CD45’s well-established role in regulating immune signaling, its therapeutic potential as an immunomodulatory target, particularly via spatial modulation or engineered sequestration, has remained largely unexplored.

In the present study, we investigate the effects of induced CD45 segregation and explore the therapeutic potential of this approach. Inspired by emerging principles in receptor clustering and agonism, we hypothesized that multivalent ligands capable of crosslinking CD45 into dense, non-signaling membrane microdomains could rewire local phosphatase accessibility and lower the TCR signaling threshold required for activation. Previous approaches to induce CD45 sequestration through receptor crosslinking have primarily relied on conventional biochemical tools such as plate-bound antibodies, polyclonal secondary antibodies, and streptavidin-biotin interactions^14,18–20^. While these studies provide consistent evidence that CD45 clustering can effectively modulate immune cell activity, they lack the precision and translational potential required for therapeutic application.

To overcome these limitations, we employed rational protein engineering strategies to design a multivalent receptor crosslinking system for precise modulation of CD45 proximity, enabling systematic exploration of immune cell signaling and function. We applied this CD45-sequestering system in preclinical murine tumor models and demonstrated that CD45 sequestration elicits curative antitumor immune responses when combined with a clinically relevant intratumorally anchored IL-12 therapy^21^. Using flow cytometry and single-cell RNA sequencing, we show that CD45 sequestration lowers the activation threshold of T cells and elicits more robust antitumor T cell responses. Our work establishes CD45 sequestration as a potential immunomodulatory strategy and provides a rationale for next-generation cancer immunotherapies that tune the spatial organization of key phosphatase signaling regulators.

## Results

### Engineering CD45 sequestering agents

Our objective is to sequester CD45 through induced receptor crosslinking, in order to attenuate its phosphatase activity (**Fig. 1 a-b**). Although mechanistically distinct from clustering-based immunoreceptor agonism, these applications share common design principles for inducing receptor clustering. Chief among them is the use of multivalent protein binders, which engage multiple non-overlapping epitopes on a single receptor, thereby promoting higher-order oligomerization and receptor crosslinking. Multiparatopic antibodies targeting two or more non-overlapping regions of cell surface receptors have been widely explored across various biological contexts, particularly in the development of immune receptor agonists and agents to induce receptor internalization or downregulation^22–26^. We based our constructs on two pan-CD45 binding clones with distinct binding epitopes, denoted αCD45 VHH and αCD45.2 mAb (**Extended Data Fig. 1a-c**)^27,28^. While CD45 is expressed in 6 different isoforms with varied extracellular domain lengths, we elected to target CD45 in an isoform-agnostic manner, due to prior evidence that complete sequestration of all isoforms is required to elicit maximal receptor sequestration^18^. As an initial approach, we engineered biparatopic fusions of these clones to a murine IgG2c Fc backbone with effector-attenuating LALA-PG mutations (**Extended Data Fig. 1c-d**). These biparatopic constructs failed to induce receptor clustering or alter downstream T cell receptor signaling, likely due to steric hindrance of these epitopes on cell-surface expressed CD45 (**Extended Data Fig. 1e-f**)^29^.

**Figure 1.**
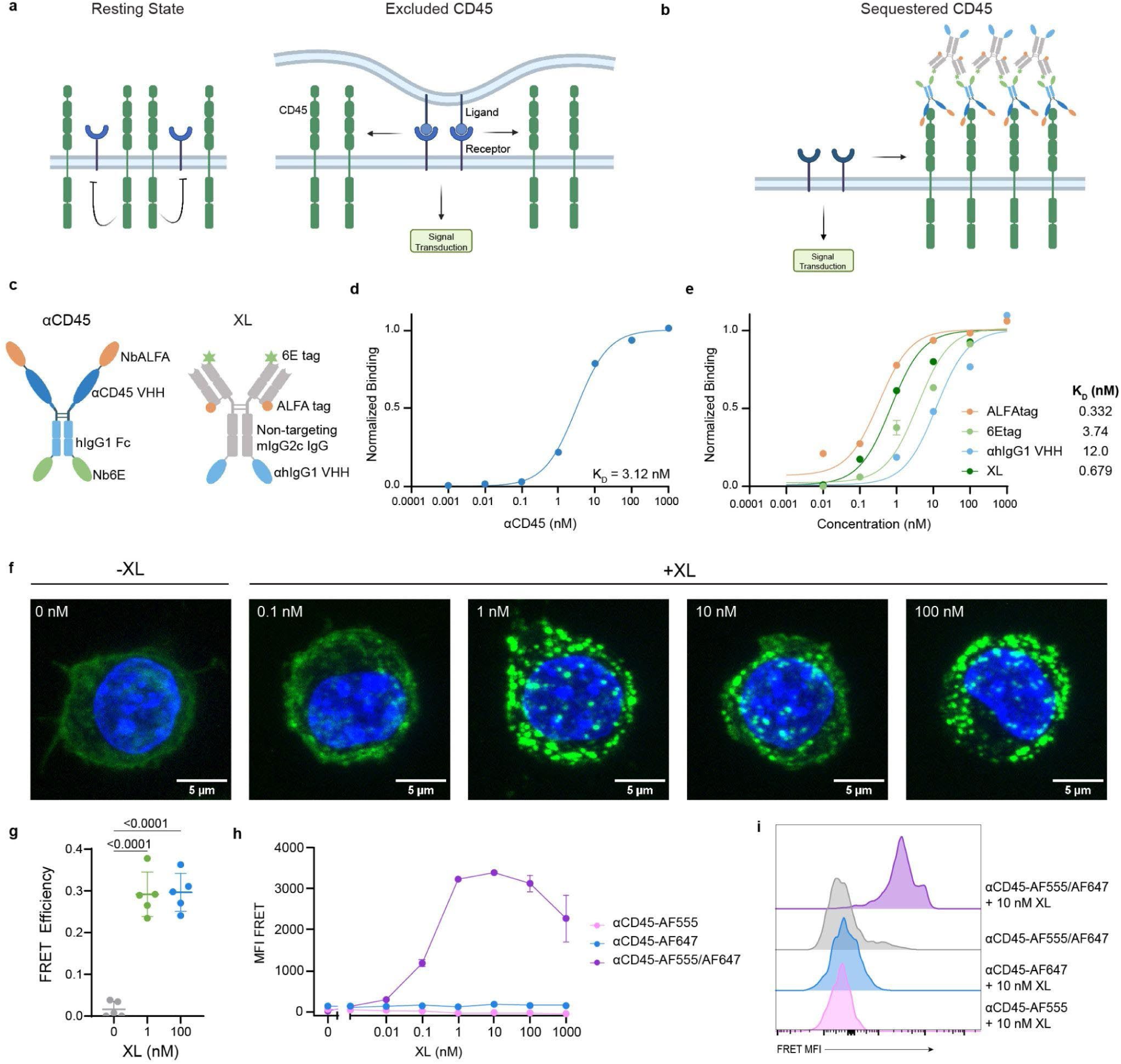
Engineered multivalent binding constructs can cluster CD45 into discrete punctae. **a.** Graphical representation of CD45 exclusion driving signal transduction at local membrane sites of ligand-receptor engagement. **b.** Graphical representation of multivalent construct induced CD45 clustering. **c.** Schematic of αCD45 and XL constructs with respective nanobody or peptide tag fusions. **d.** Equilibrium binding curves normalized to B_max_ of aCD45 on murine splenocytes (n=3 ± s.d.). Binding was measured by AF647 MFI and normalized to the Bmax values. **e.** Equilibrium binding curves normalized to B_max_ for individual peptide tags or XL construct on murine splenocytes pulsed with aCD45 for 30 min. (n=3 ± s.d.). Equilibrium dissociation constants (K_D_) in **d-e** were calculated using a nonlinear regression fit for one-site total binding with no nonspecificity. **f.** Confocal Z-stack projections of RAW264.7 CD45-mGreenLantern pulsed with 100 nM αCD45 for 30 min and treated with titrations of XL. CD45 is shown in green and nuclei in blue. **g.** FRET efficiency calculated using the FRET acceptor photobleaching method on a Leica SP8 confocal with a 63x oil objective (mean ± SD; n = 5). *p* values were determined by one-way ANOVA followed by Tukey’s multiple-comparison test. **h.** FRET MFI on murine splenocytes, measured in the PE-Cy5 channel (561 nM excitation laser, 670/30 nM emission filter) via flow cytometry (mean ± SD; n = 3). **i.** Select representative histograms of FRET signal.

Given the lack of published CD45 binders targeting additional non-sterically hindered CD45 epitopes at the time, we sought an alternative protein engineering strategy using a two-component system. The first element of this approach is a binder that directly engages CD45, hereafter referred to as “αCD45”, while the second component induces the cross-linking of the CD45 binder, and is hereafter referred to as crosslinker, or “XL” (**Fig. 1c**). To prepare the αCD45 construct we fused the αCD45 VHH to a human IgG1 Fc domain containing effector-attenuating LALA-PG mutations. To increase the number of epitopes available for crosslinking, we identified two short 14-16 amino acid peptide tags with corresponding anti-tag nanobodies (termed the ALFA and 6E binding pairs) from the literature^30,31^. We conjugated the ⍺ALFA and ⍺6E VHHs to the N-and C-termini of the ⍺CD45 construct, respectively, to produce a final antibody that could recognize both CD45 as well as the ALFA and 6E peptides. To prepare the highly multivalent “XL” construct, we fused the corresponding ALFAtag, 6Etag, as well as an αhIgG1-specific VHH to the respective termini of an irrelevant fluorescein-specific antibody on a murine IgG2c LALA-PG backbone, such that the XL antibody can propagate crosslinks from six different sites (**Extended Data Fig. 2a**)^32^. We expressed and validated these constructs, confirming that the artificial epitopes on αCD45 were sterically accessible when bound to surface-expressed CD45 (**Fig. 1d-e, Extended Data Fig. 2b-d**).“

To evaluate whether the engineered αCD45 and XL system was capable of inducing higher order receptor crosslinking of CD45, we utilized RAW264.7 cells stably expressing CD45-mGreenLantern. Cells first pulsed with αCD45, followed by treatment with increasing concentrations of XL, resulted in the formation of discrete CD45 puncta visualized via confocal microscopy (**Fig. 1f**). We further confirmed via Förster Resonance Energy Transfer (FRET) acceptor photobleaching that stepwise incubation of αCD45-AF555 and αCD45-AF647, followed by XL resulted in CD45 clustering on the cell surface (**Fig. 1g**). Using flow cytometry to more precisely quantify differences in FRET efficiency, we observe a dose-dependent increase in FRET signal correlating with XL concentration, confirming the robust formation of dense CD45 receptor aggregates (**Fig. 1h-i**). Notably, the αCD45 + XL system is capable of maintaining CD45 crosslinking on the cell surface for up to 24 hours without significant internalization of the receptor (**Extended Data Fig. 2e-g**)^33,34^. The XL design shown was selected through a systematic and extensive evaluation of artificial epitope orientation and valency on the IgG scaffold, identifying the configuration that elicited maximal CD45 receptor crosslinking. We found that the efficiency of CD45 clustering was dictated by the orientation of the ahIgG1 VHH fusion terminus (**Extended Data Fig. 3a-d**). Triepitopic XL constructs clustered CD45 and activated T cells more efficiently than biepitopic designs (**Extended Data Fig. 4a-d**). By engineering an artificial multivalent high-affinity interface onto the CD45 extracellular domain, we achieved precise, tunable control of its local proximity and spatial organization.

### CD45 sequestration lowers the signaling threshold in T cells

We sought to characterize the effects of altered CD45 localization on TCR signaling. CD45 sequestration increased phosphorylation at both Y394 (activating site) and Y505 (inhibitory site) on Lck, and enhanced phosphorylation of the CD3ζ chain at Y142, a downstream marker of TCR engagement (**Fig. 2a-c**). CD45 sequestration also increased global phosphotyrosine abundance across the T cell proteome, demonstrating an overall enhancement in basal T cell signaling activity (**Fig. 2d**).

**Figure 2.**
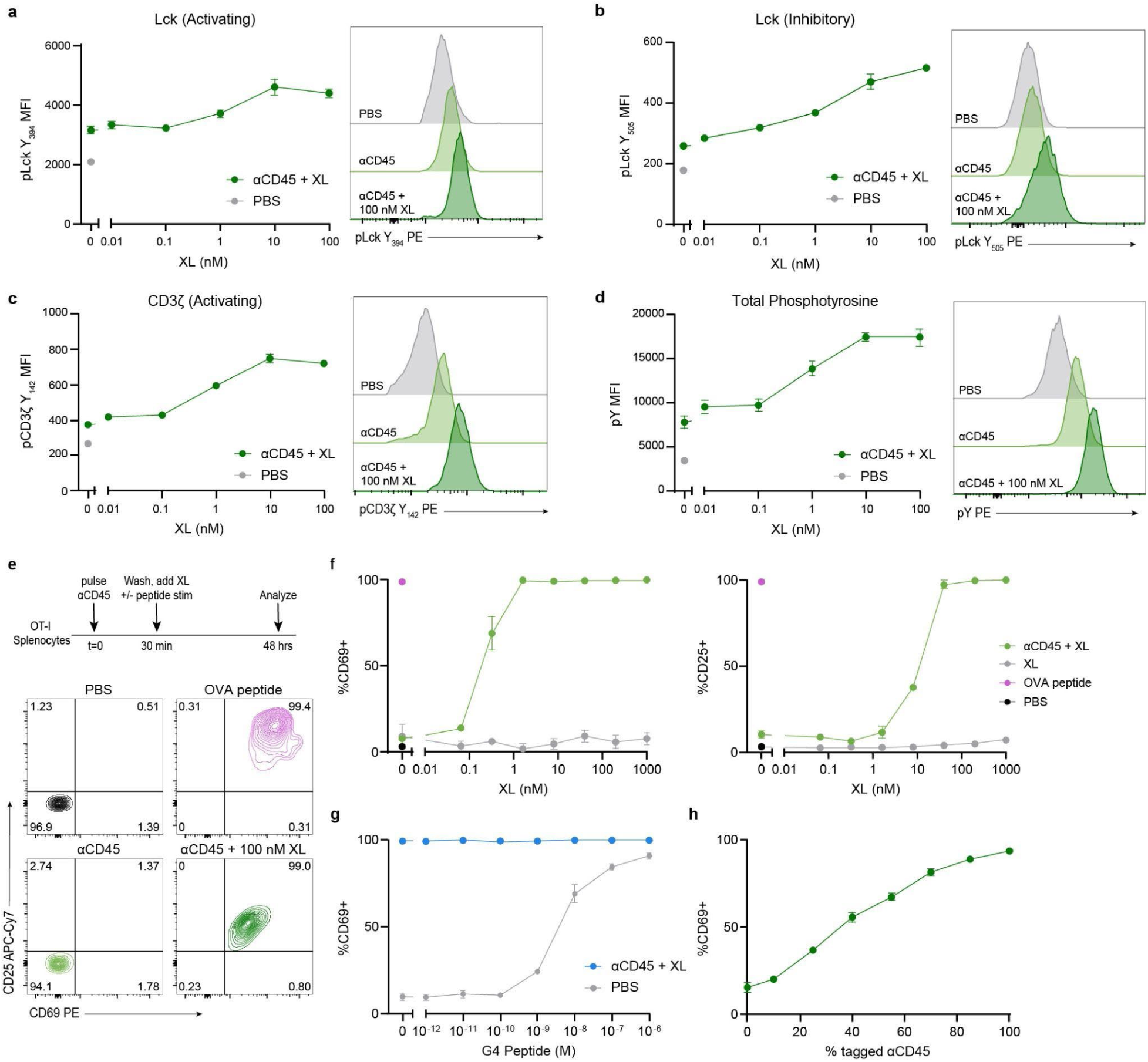
CD45 sequestration lowers the TCR signaling threshold and activates T cells. a-**d.** Primary CD8^+^ T cells were pulsed with αCD45 and treated with titrations of XL. Cells stained for flow cytometry analysis of phosphorylation motifs on Lck Y394 (**a**), Lck Y505 (**b**), CD3z Y142 (**c**), and total phosphotyrosine (**d**) (mean ± SD; n = 3). **e.** Experimental setup for OT-I T cell activation assay in **f-h** and representative CD69 and CD25 gating. **f.** Percent CD69 and CD25+ OT-I T cells. OT-I splenocytes were pulsed with 100 nM αCD45, washed twice and treated with titrations of XL (mean ± SD; n = 3). **g.** Percent CD69+ OT-I T cells. OT-I splenocytes were pulsed with 100 nM αCD45, diluted by indicated percentages of single domain αCD45 VHH, washed twice and treated with 100 nM of XL (mean ± SD; n = 3). **e.** Percent CD69+ OT-I T cells. OT-I splenocytes were treated with titrations of G4 peptide, in the presence or absence of αCD45 + XL. Cells were pulsed with 100 nM αCD45, washed twice and treated with 100 nM of XL (mean ± SD; n = 3).

To evaluate if the magnitude of these CD45 sequestration-driven phosphorylation changes could alter T cell activation states, we treated splenocytes isolated from OT-I transgenic mice with αCD45 and XL. 48 hours post-treatment, we observed strong upregulation of canonical T cell activation markers CD69 and CD25 (**Fig. 2e-f**). Interestingly, CD45 sequestration alone was sufficient to activate T cells, irrespective of TCR stimulation strength, evidenced by the addition of a low-affinity ovalbumin-derived peptide (G4; SIIGFEKL) (**Fig. 2g**). To determine the extent of CD45 engagement required for maximal T cell activation, we performed a competitive titration using αCD45 diluted with the unconjugated αCD45 VHH domain. We observed a strong linear relationship between CD69 expression and the proportion of sequestered CD45, indicating that full sequestration is required for maximal activation, but even partial displacement of CD45’s local proximity is sufficient to alter TCR phosphorylation dynamics (**Fig. 2h**). Thus, CD45 sequestration can lower the TCR signaling threshold and activate T cells by reprogramming the local phosphatase landscape.

### CD45 sequestration synergizes with local IL-12

To assess whether the T cell activation observed in vitro with CD45 sequestering constructs could be recapitulated *in vivo*, we sought to evaluate the effects of CD45 sequestration in murine syngeneic tumor models. We selected a dose of αCD45 sufficient to achieve saturating receptor occupancy across all immune cell subsets and tissue compartments (**Extended Data Fig. 5a-e**) and administered an equimolar dose of the XL.

To ensure adequate receptor engagement and clearance of unbound αCD45 from circulation, we administered αCD45 intratumorally 24 hours prior to intratumoral injection of the XL. We tested this therapeutic regimen (denoted “CD45 XL”) in two widely used syngeneic tumor models: MC38 colorectal carcinoma, which is partially responsive to checkpoint blockade immunotherapy (CBI), and B16F10, an aggressive melanoma model with poor responsiveness to CBI. CD45 sequestration alone did not confer a measurable survival benefit, but was well tolerated, with no observed weight loss or signs of treatment-related toxicity (**Extended Data Fig. 6a-d**).

Although CD45 sequestration monotherapy confers no survival benefit, we reasoned that it may amplify T cell priming induced by other immunostimulatory agents. We sought to evaluate the therapeutic potential of CD45 sequestration in combination with interleukin 12 (IL-12). We have previously reported a locally retained form of IL-12 conjugated to aluminum hydroxide, which significantly mitigated systemic toxicity while eliciting potent antitumor responses when used in combination with PD-1 checkpoint blockade^35^. This locally anchored IL-12 formulation is now advancing in Phase 1 clinical trials^21^. Two days following IL-12 administration, timed to coincide with peak tumor neoantigen uptake in the tumor-draining lymph node (tdLN)^35^, we administered CD45-sequestering constructs. A second dose of CD45-sequestration therapy was repeated four days later. Strikingly, the combination of local IL-12 and CD45 XL treatment elicited synergistic tumor growth delay in the B16F10 syngeneic melanoma model, as compared to local IL-12 alone (Bliss independence test, *p* = 7.41e^-5^) (**Fig. 3a-b**). Animals experienced no treatment related toxicity, as assessed by weight loss (**Fig. 3c**). We confirmed the synergy between CD45 sequestration and IL-12 treatment in the more immunogenic MC38 model, where IL-12 monotherapy delayed tumor growth, but its combination with CD45 XL resulted in complete tumor rejection in 100% of treated mice (**Fig. 3d-e**). Co-administration of IL-12 with either αCD45 or XL alone elicited no survival benefit over IL-12 monotherapy, demonstrating that CD45 clustering, and therefore sequestration is required for tumor growth delay (**Fig. 3f**). These findings highlight the potential of CD45 sequestration as a novel therapeutic strategy that enhances the efficacy of localized cytokine therapy.

**Figure 3.**
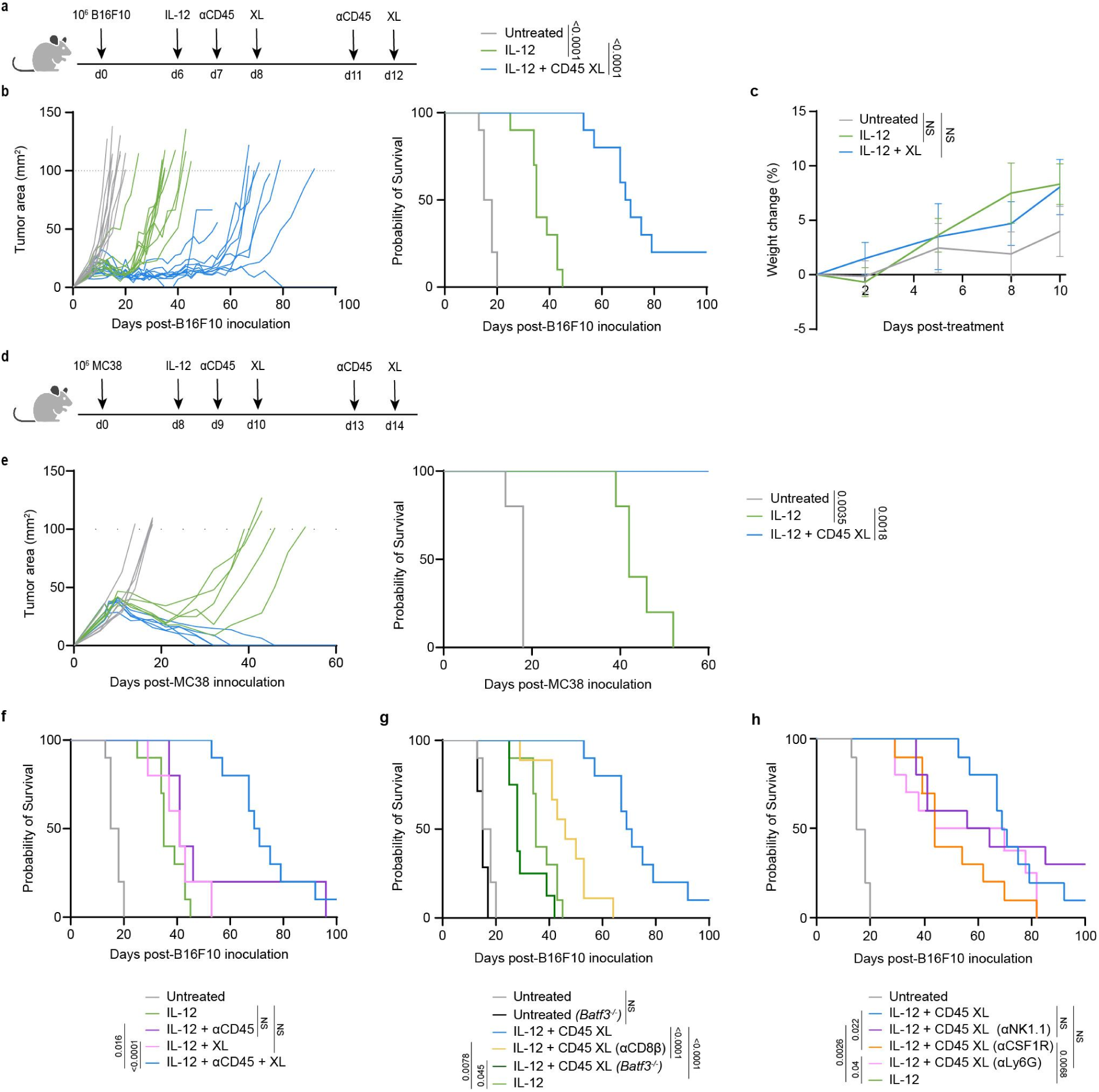
CD45 sequestration synergizes with intratumorally anchored IL-12 therapy. **a.** Mice (*n* =10/group) bearing B16F10 tumors treated with 5 ug ABP tagged IL-12 and 100 µg of alum, 40 µg of aCD45 and 60 µg of XL as shown. **b.** (left) B16F10 tumor growth curves and (right) Kaplan Meier survival. **c.** Percent weight change measured from the start of treatment on day 6 (n = 5/group). **d.** Mice (*n* =5/group) bearing MC38 tumors treated with 5 ug ABP tagged IL-12 and 100 µg of alum, 40 µg of aCD45 and 60 µg of XL as shown. **e.** (left) MC38 tumor growth curves and (right) Kaplan Meier survival. **f.** Treatment regimen shown in **a**, administered with either αCD45, XL, or both agents. **g-h.** Treatment regimen shown in **a**, in combination with depleting antibodies or performed in Batf3 knockout mice.

### Combination therapy promotes tumor-reactive CD8**^+^** T cell responses

To interrogate the immune subsets responsible for tumor growth delay in the IL-12 + CD45 XL combination therapy, we evaluated our treatment paradigm using various knockout mice and depleting antibodies. The efficacy of CD45 XL combination therapy was dependent on CD8^+^ T cells and Batf3^+^ dendritic cells, underscoring the essential role of tumor antigen cross-presentation and CD8^+^ T cell priming (**Fig. 3g**). NK cells, neutrophils, and monocytes were found to be dispensable for anti-tumor efficacy (**Fig. 3h**). We observed modest survival contributions from CSF1R^+^ macrophages, which may reflect a supportive role for macrophages in antigen presentation and T cell priming (**Fig. 3h**)^36,37^.

Given the requirement for CD8^+^ T cells, we next sought to dissect how IL-12 and CD45 sequestration synergized to drive tumor control, focusing on the generation of tumor-specific CD8^+^ T cell responses as the lynchpin in productive antitumor immune responses. We administered the full IL-12 + CD45 XL regimen to B16F10 tumor-bearing mice, and two days after the final XL dose, profiled tissues for tumor-reactive T cell responses (**Fig. 4a**). To assess these responses, we treated B16F10 tumor-bearing mice and performed interferon gamma (IFNγ) ELISPOTs on splenocytes two days after the final XL dose. The combination treatment with CD45 XL elicited a significant increase in the tumor-specific responses relative to IL-12 monotherapy (**Fig. 4b**). robust T cell priming and activation within secondary lymphoid organs has been widely recognized as essential for effective immune-mediated tumor control^38,39^. To evaluate the phenotypic features underlying the effects of IL-12 + CD45 XL combination therapy on tumor-specific CD8⁺ T cells, we focused on T cells reactive to the immunodominant p15E endogenous retroviral antigen expressed by B16F10 tumors^40,41^. In the tumor-draining lymph node (tdLN), we observed a higher frequency of p15E tetramer-reactive CD8⁺ T cells following combination therapy, relative to IL-12 monotherapy (**Fig 4c-d**). While the absolute number of p15E-reactive CD8^+^ T cells was not elevated relative to IL-12 monotherapy in the tdLN (**Fig. 4e**), we saw striking differences in the phenotypic states of these T cells. Within the tdLN, p15E^+^ T cells from mice treated with CD45 XL combination therapy exhibited enhanced cytolytic potential, evidenced by a significant increase in the proportion and absolute count of the Granzyme B positive (GzmB^+^) p15E^+^ T cell population (**Fig. 4f-h**). p15E-reactive CD8^+^ T cells treated with CD45 XL exhibited enhanced activation states, and expansion of the CD25^+^ compartment (**Fig. 4i-j**). In contrast, the phenotype of p15E-reactive T cells in the spleen did not differ between monotherapy and combination treatment (**Extended Data Fig. 7a-d)**. CD45 sequestration was also not the dominant factor in promoting increased infiltration in the tumor, suggesting that its primary mechanism of action is on newly primed T cells in the tdLN (**Extended Data Fig. 7e-g**).

**Figure 4.**
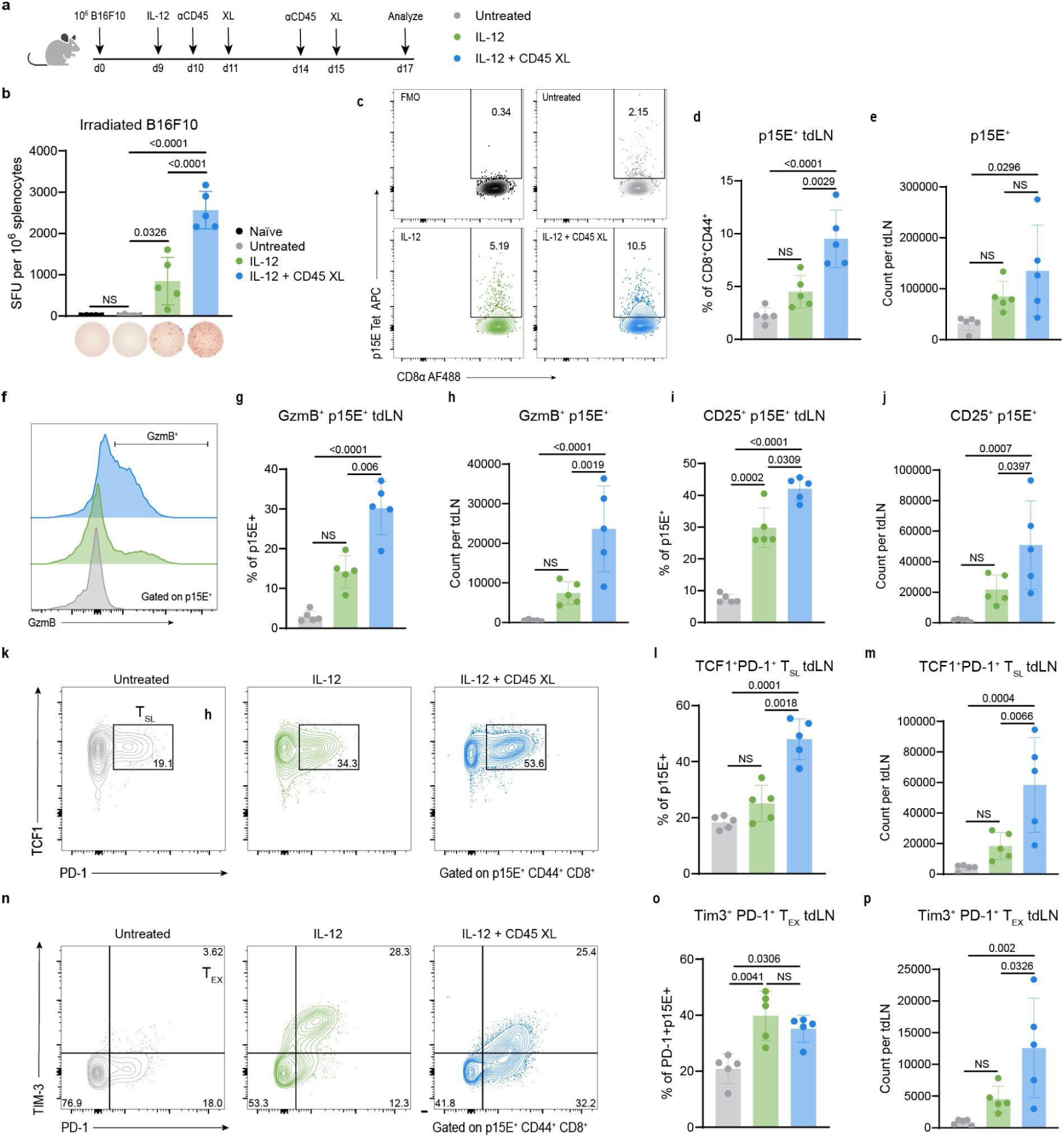
CD45 sequestration combination therapy promotes a tumor-reactive T cell response in the tdLN. **a.** C57BL/6 mice (n=5) were inoculated with 1×10^6^ B16F10 cells and treated as shown. Tissues were harvested for flow cytometry or ELISPOT analysis 48 hours after the completion of treatment. **b.** Number of spot-forming units (SFU) per 10^6^ splenocytes (mean ± s.d., n=5/group) in an IFN-γ ELISPOT assay, and representative images of ELISPOT wells. **c.** Representative contour plots and gating for p15E tetramer-reactive cells in the tdLNs, previously gated on live CD8^+^CD44^+^ cells. Treatment effects on the **d.** proportion and **e.** count of p15E^+^ tumor-reactive cells in tdLNs. **f.** Representative histogram of granzyme B (GzmB) expression in p15E^+^CD8^+^ T cells. Treatment effects on the **g.** proportion and **h.** count of GzmB^+^ p15E^+^ tumor-reactive cells in tdLNs. Treatment effects on the **i.** proportion and **j.** count of CD25^+^ p15E^+^ tumor-reactive cells in tdLNs. **k.** Representative stem-like T cell (Tcf1^+^PD-1^+^) gating. tdLN proportion (**l**) and count (**m**) of stem-like PD-1^+^TCF1^+^ p15E^+^CD8^+^ T cells. **n.** Representative exhausted T cell (PD-1^+^TIM3^+^) gating. tdLN **o.** frequency and **p.** count of exhausted PD-1^+^TIM3^+^ p15E^+^ T cells. *p* values were determined by one-way ANOVA followed by Tukey’s multiple-comparison test.

Recent work has highlighted the importance of TCF1^+^ PD-1^+^ stem-like T cells residing in the draining lymph nodes at disease sites as a critical reservoir for sustained effector cell generation in the context of cancer and viral infections^42–45^. The robust tumor-reactive T cell activation observed in treated tdLNs prompted us to investigate the impact of CD45 sequestration combination therapy on the magnitude of the stem-like compartment (T_SL_, TCF1^+^PD-1^+^). Combination therapy led to a significant increase in the relative proportion of stem-like T cells compared to IL-12 alone, suggesting that CD45 sequestration supports the maintenance and expansion of this progenitor population during effector differentiation (**Fig. 4k-m**). Encouragingly, combination therapy did not drive further differentiation into an exhausted phenotype (T_EX_, Tim3^+^PD-1^+^), as the frequency of exhausted effectors, marked by Tim3 expression, remained equivalent between monotherapy and combination groups (**Fig. 4n**). Notably, combination therapy induced a significantly greater expansion of the exhausted T cell compartment by absolute count relative to IL-12 monotherapy, while maintaining the stem-like reservoir (**Fig. 4o**). We observed similar responses at the total CD8+ T cell compartment level, with CD45 sequestration combination therapy leading to an increase in the proportion, activation state and stem-like phenotypes of effector/effector memory (Eff/EM, CD44^+^CD62L^-^) CD8^+^ T cells in the tdLN, compared to IL-12 monotherapy (**Extended Data Fig. 8a-o**). Collectively, these results suggest that CD45 sequestration amplifies T cell activation and differentiation, while maintaining the progenitor reservoir necessary for sustained antitumor immunity.

### CD45 sequestration broadens T cell clonotypes

To further evaluate the clonotypic and phenotypic features underlying the effects of IL-12 + CD45 XL combination therapy, we profiled the intratumoral p15E^+^ tumor-reactive CD8⁺ T cells using single-cell RNA (scRNA) and TCR sequencing (**Fig. 5a**). After filtering out naïve-like and low-quality cells, dimensionality reduction via UMAP and Louvain clustering revealed five transcriptionally distinct states: precursor-exhausted, progenitor-exhausted, proliferating, terminally exhausted, and KLR⁺ exhausted T cells (**Fig. 5b-d**), consistent with established T cell atlases^38^. Precursor and progenitor-exhausted T cells expressed high levels of stemness genes such as *Tcf7*, *Bach2*, and *Stat1*; proliferating cells upregulated the replication markers *Mki67*, *Cenpe*, and *Top2a*; and terminally exhausted cells expressed classical exhaustion markers (*Lag3*, *Tigit*, *Havcr2*, *Slamf7*), with downregulation of *Tcf7* (**Fig. 5d, Extended Data Fig. 9a, c**). KLR exhausted cells additionally upregulated *Klrg1*, *Klrd1*, and *Klrc1* (**Fig. 5d, Extended Data Fig., 9b, c**).

**Figure 5.**
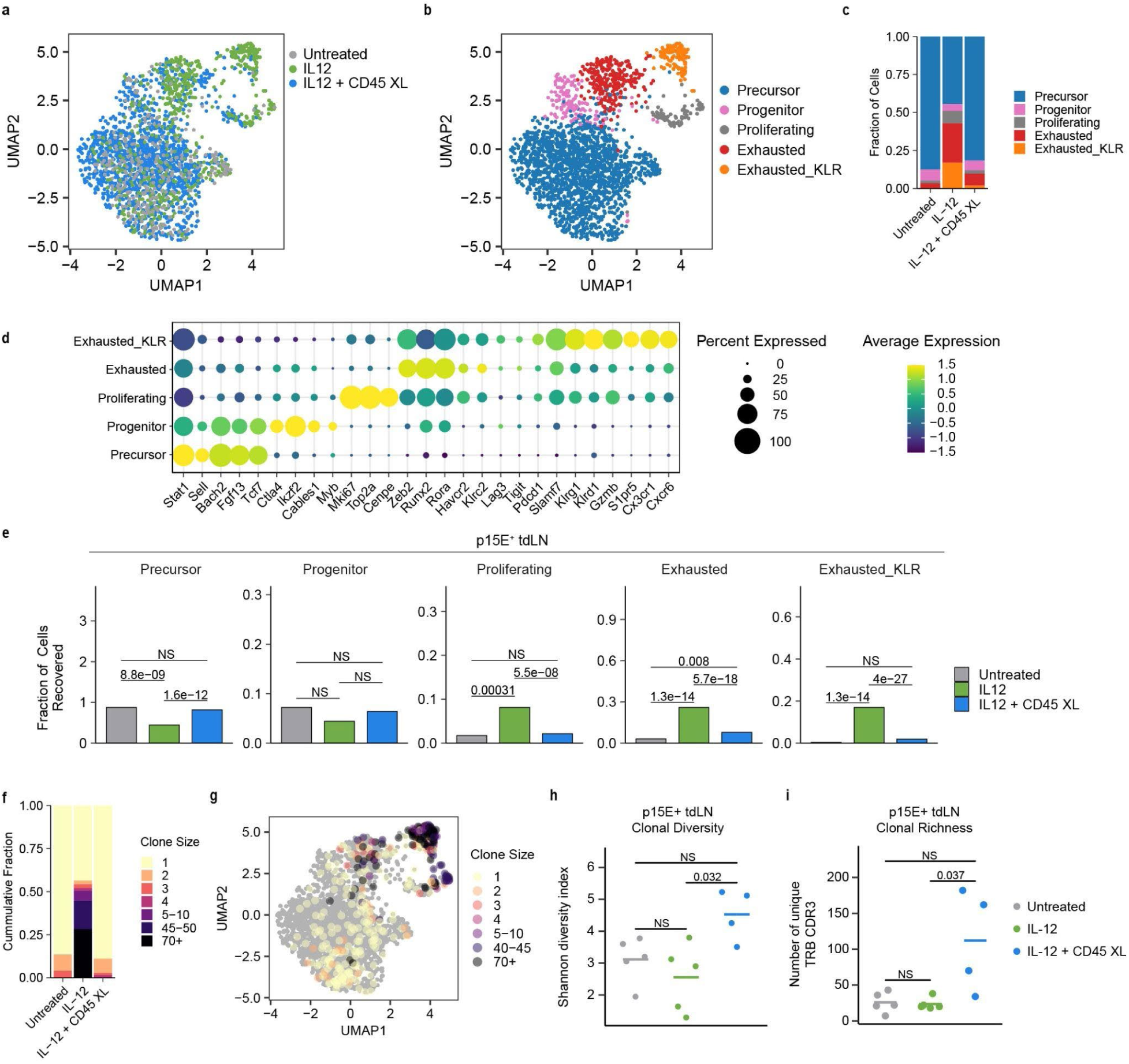
CD45 sequestration combination therapy sustains a tumor-specific T cell response and broadens the tumor-reactive clonal repertoire in the tdLN. Mice bearing B16F10 tumors (n = 4-5/group) were harvested 48 h. after the completion of treatment and sorted for CD8^+^ T cells specific for the immunodominant p15E retroviral antigen in the tdLN for downstream scRNA-seq. **a.** UMAP of p15E^+^ CD8^+^ T cells in the tdLN, by treatment group. **b.** UMAP of sequenced T cells, by phenotype. **c.** Frequency of p15E^+^ T cell phenotypes recovered from each treatment group. **d.** Bubble plot of scaled gene expression of marker genes in p15E^+^ T cell phenotypes. **e.** Frequency of cellular transcriptional states in the p15E^+^ tdLN by treatment group. p values were calculated using Fisher’s exact test for significant proportions. **f.** Stacked bar plots of clonal sizes among tdLNs, of untreated, IL-12 or IL-12 + CD45 XL mice. **g.** UMAP of p15E^+^ T cells, colored by clone size. **h.** Shannon diversity of TCR repertoire in p15E^+^ tdLN. **i.** Clonal richness of TCR repertoire in p15E^+^ tdLN. p values (**h-i**) were calculated using a two-sided Wilcoxon rank sum test and adjusted Bonferroni correction.

We next examined the frequency of each transcriptional phenotype among the recovered p15E-reactive CD8^+^ T cells across each treatment group (**Fig. 5c, e**). A significantly higher proportion of the cells recovered in the combination treatment were found to reside in the precursor-exhaustion state, and retained stem-like transcriptional signatures, compared to IL-12 monotherapy (**Fig 5c, Extended Data Fig. 10a-b**). Conversely, the IL-12 monotherapy treatment exhibited a significantly higher proportion of terminally exhausted T cell states (**Fig. 5e**). Consistent with the enrichment of stem-like progenitor CD8⁺ T cells seen via flow cytometry (**Fig. 4k-m**), these results suggest that CD45 sequestration combination therapy limits the progression of exhaustion and supports a more durable, functional T cell pool.

Given CD45’s established role in antigen discrimination and suppression of low-avidity T cell responses, we next assessed how CD45 sequestration impacts newly primed TCR clonotype diversity^16,17^. To determine the effect of CD45 sequestration on newly primed TCR clonotypes, we performed TCR repertoire sequencing on the p15E-reactive CD8^+^ T cells in the TDLN (**Fig. 5f-i**). TCR sequencing of p15E-reactive CD8⁺ T cells revealed increased clonal diversity and a greater clonal richness (i.e. total number of p15E-reactive clonotypes) in the combination group, consistent with enhanced priming of a broader TCR repertoire (**Fig 5h-i**). These clones exhibited reduced clonal expansion under combination therapy, fostering a more diverse tumor-reactive immune response (**Fig.5f**). Taken together, our findings demonstrate that CD45 sequestration synergizes with IL-12 to enhance the magnitude, diversity and functional quality of tumor-specific CD8⁺ T cell responses, while limiting terminal exhaustion and preserving stem-like potential.

## Discussion

This study offers new insights into harnessing CD45 localization as an effective strategy to modulate T cell activation and boost antitumor immunity. Although CD45 has been leveraged as a therapeutic target, either by exploiting its phosphatase activity to regulate receptor signaling^46,47^ or as an anchoring site for targeted delivery to lymphocytes,^33,34^ modulating its activity through kinetic segregation remains largely unexplored. Using rational protein engineering and established design principles for receptor clustering, we developed a tunable system to control CD45 localization at the cell surface. This modular approach provides precise control over valency, affinity, and spatial arrangement of receptors, enabling rapid *in vivo* screening of receptor clustering without the need for bespoke noncompetitive binders. Using this modular system, we show that induced CD45 segregation rewires the local phosphatase landscape and lowers the TCR activation threshold, enabling T cell responses to otherwise weak or subthreshold stimuli. CD45 sequestration alone was insufficient to initiate a productive antitumor response, presumably due to an insufficient baseline level of TCR signaling in secondary lymphoid organs to generate productive tumor-reactive T cell responses.

Our findings demonstrate that CD45 sequestration can synergize with a localized cytokine therapy to enhance the generation of activated effector T cells in the tdLN and drive curative antitumor immune responses. Encouragingly, we saw no signs of toxicity associated with CD45 modulation as assessed by treatment associated weight loss. Consistent with CD45 sequestration lowering the activation threshold for T cells, we observe broader clonotypic expansion of tumor-reactive T cells following antigen generation by local IL-12 treatment. The emergence of new clonotypes in the combination therapy suggests that these may be low-affinity TCRs, previously non-functional without the addition of CD45 sequestration. Notably, low-affinity T cells have been shown to retain stem-like properties (as shown in Figure 5), maintain effector functions, and exhibit superior tumor control, due to reduced chronic TCR signaling and exhaustion in antigen rich environments^48–51^.

IL-12 served as an effective proof-of-concept for demonstrating the therapeutic potential of CD45 sequestration, but it represents only one of many possible immunomodulatory stimuli. We anticipate CD45 sequestration might prove effective in combination with treatments that stimulate antigen release and cross-priming, such as radiation therapy^52^, neoantigen vaccination strategies, or other cytokine therapies. Such combinations may further improve the breadth and magnitude of anti-tumor immune responses and help overcome resistance in poorly immunogenic tumor settings.

In conclusion, our results provide compelling evidence that CD45 spatial reorganization is not merely a passive consequence of immunological synapse formation, but rather a tunable axis of immune modulation. These findings position CD45 sequestration as a tractable and novel paradigm for modulating immune signaling, with potentially broad applications in immunotherapy.

## Methods

### Cell lines

B16F10 (ATCC), MC38 (a gift from J. Schlom, National Cancer Institute), and RAW264.7 (ATCC) cell lines were cultured in Dulbecco’s Modified Eagle’s Medium (DMEM) (ATCC) supplemented with 10% FBS and 100 U/mL Penicillin-Streptomycin (Gibco). Adherent cells were cultured at 37°C and 5% CO_2_. Expi293F cells (Gibco) were cultured in Expi293 Expression Medium (Gibco), in baffled flasks shaking at 37 °C and 8% CO_2_. All cell lines were validated as pathogen-free via IMPACT and mycoplasma testing.

### Mice

Female C57BL/6J mice were purchased from Jackson Labs (000664). OT-I mice (C57BL/6-Tg(TcraTcrb)1100Mjb/J) were a generous gift from Darrell Irvine and maintained in-house. Batf3^-/-^ mice (B6.129S(C)-Batf3tm1Kmm/J) were a generous gift from Stefani Spranger and bred and maintained in-house. Age matched 6–10-week-old females were used in experiments. All animal studies and procedures were conducted under the approval of the Massachusetts Institute of Technology Committee on Animal Care in accordance with federal, state, and local guidelines.

### Cloning and protein purification

Recombinantly expressed proteins were cloned into the gWiz vector (Genlantis) using InFusion cloning (Takara Bio) or using the NEBridge Golden Gate Assembly Kit BsaI-HF v2 (NEB), according to the manufacturer’s protocols. Fam20C kinase and IL12-ABP were previously cloned into the gWiz vector^35^. Plasmid sequences were confirmed by Sanger sequencing (Quintara Biosciences) or nanopore sequencing (Plasmidsaurus), transformed into Stellar Competent Cells (Takara Bio) and purified using the NucleoBond Xtra Midi endotoxin-free midi-prep kit (Takara Bio). Amino acid sequences are listed in Supplementary Table 1.

Protocols for recombinant protein production in Expi293F cells (Gibco) were adapted from Zhou et. al., 2023^53^. Briefly, 8 mg/L of polyethylenimine (Polysciences 23966) prediluted in Opti-MEM medium (Gibco) was mixed with 1 mg/L plasmid DNA in a total volume of 125 mL/L Opti-MEM. For full length IgG production, a 1:1 mass ratio of heavy and light chain plasmid was used. For alum anchored IL-12, a 9:1 mass ratio of IL-12-ABP plasmid and Fam20C plasmid was used. After a 15 min incubation at room temperature, DNA-PEI complexes were added dropwise to 3 x 10^6^ cells/mL of Expi293F culture. 24 hrs. post-transfection, 25 mL/L of 20% w/v Soy C-CELL S146B Peptone (Organotechnie) sterile filtered in Expi293F Media (Gibco) and 8 mL/L of 500 mM valproic acid (Sigma) sterile filtered in PBS (Corning) were added to the culture. 6 days after transfection, proteins were purified from cell culture supernatants using Ni Sepharose High Performance Resin (Cytiva) for 6xHis tagged proteins, and rProtein A Sepharose Fast Flow Resin (Cytiva) for proteins containing an IgG Fc domain. All recombinant proteins were validated for size by SDS-PAGE. Proteins were run alongside the Novex Sharp Pre-Stained Protein Standard (Invitrogen) on a NuPAGE 4 to 12% Bis-Tris gel (Invitrogen) with 2-(N-morpholino) ethanesulfonic acid (MES) running buffer and stained for visualization with SimplyBlue Safe Stain (Life Technologies). Proteins were further purified by size exclusion chromatography via a Superdex 200 Increase 10/300 GL column (Cytiva) on an AKTA FPLC system (Cytiva, formerly GE Healthcare). Proteins were buffer exchanged into PBS using appropriate Amicon spin columns (Milipore Sigma). Purified proteins were confirmed to have low endotoxin levels (<0.1 EU per dose) by the Endosafe^TM^ Nexgen-PTS system (Charles River).

### ELISA and cell binding assays

Clear Flat-bottom Immuno Nonsterile Nunc 96-well MaxiSorp Plates (Invitrogen) were coated overnight at 4 °C in PBS (Corning). Wells were washed 3 times with PBST (PBS supplemented with 0.05% v/v Tween-20 (Millipore-Sigma)). Blocking was performed in PBSTA (1% w/v BSA (Sigma Aldrich) and 0.05% v/v Tween-20 in PBS) overnight. Subsequent binding incubations were performed for 30 min-2 h at RT in PBSTA, and wells were washed 4 times with PBST in between incubations. Detection was performed via HRP-conjugated anti-mouse IgG secondary diluted 1:10,000 (Abcam) or HRP-conjugated streptavidin diluted 1:15,000 (ThermoFisher). 1-Step ultra TMB-ELISA substrate solution (ThermoFisher) was added to develop for 5-10 minutes and quenched with 2N sulfuric acid (Sigma). Absorbance wavelength at 450 nm and reference wavelength at 570 nm were measured on an Infinite M200 microplate reader (Tecan). Equilibrium binding constants (K_D_) were calculated using the one site, total equation in GraphPad Prism software V10.

Spleens from naïve mice were harvested and processed into single-cell suspensions through a 70-micron cell strainer (Grenier). Red blood cells were lysed using ACK Lysis Buffer (Gibco). Splenocytes were plated in 96 well TC-treated U bottom plates (Falcon) at a concentration of 100,000 cells/well in RPMI supplemented with 10% FBS (Gibco), 100 U/mL Penicillin-Streptomycin (Gibco), 1x sodium pyruvate (Thermo Fisher Scientific), 1x nonessential amino acids (Thermo Fisher Scientific) and 1x 2-mercaptoethanol (Thermo Fisher Scientific). Cells were treated with titrations of AF647 labeled proteins at the indicated concentrations for 2 h. at 4°C. AF647 MFI was quantified using a BD FACSymphony A3 and equilibrium binding constants (K_D_) were calculated using the one site, total equation in GraphPad Prism software V10.

### Generation of CD45-mGreenLantern cell lines

sgRNA’s against the C-terminal CD45 locus were designed using Benchling’s CRISPR Guide RNA Design Tool and cloned into the pSpCas9(BB)-2A-GFP vector (plasmid PX458, a generous gift from the Zhang lab)^54^. The HDR template was designed to introduce the mGreenLantern fluorescent protein at the C-terminus of CD45, with 500bp of flanking homology regions to the genome. Single stranded DNA HDR template was generated using Guide-it™ Long ssDNA Production System (Takara). 2×10^6^ RAW264.7 cells were electroporated with the Nucleofector 4D program (Lonza, program DS-36). 5 μg of px458 plasmid and 5 μg of linear HDR template were mixed in 100 μL SF buffer (Lonza) containing RAW264.7 cells. Following electroporation, cells were cultured in 10 mL complete media for 48 hrs. Cells were dissociated with CellStripper (Corning), and FACS sorted to isolate Cas9^+^ cells based on mCherry fluorescent protein expression. Cells were recovered in culture for 7 days and then FACS sorted for HDR+ cells based on mGreenLantern expression. Clonal cell populations were sorted into 96 well plates, and single clones were selected based on mGreenLantern expression on flow cytometry. Genomic DNA from single clones was extracted with the DNeasy Blood & Tissue Kit (Qiagen), and regions of the CD45 locus were PCR amplified. Genomic insertion was verified with Sanger sequencing.

### Microscopy

RAW264.7 CD45-mGreenLantern cell line was seeded overnight at 10,000 cells/well in Poly-D lysine-coated PhenoPlate™ 96-well microplates (Revvity). Cells were pulsed with 100 nM αCD45 for 30 minutes at 37°C, washed twice in media, and resuspended in 200 μL XL at the indicated concentrations. Following incubation, media was removed, and cells were washed twice with PBS before being immediately fixed with an equal volume of BD Phosflow Fixation Buffer I (BD Biosciences) at 37°C for 10 minutes. Cells were washed twice with PBS, and stained 1:20,000 with DAPI (5 mg/mL, ThermoScientific) for 5 min. at room temperature. Cells were washed twice with PBS and kept at 4°C protected from light prior to Z-stack imaging with a Leica SP8 scanning confocal microscope using a 100x silicone oil-based objective. Images were processed with ImageJ (Fiji). Imaging was performed in two or more independent replicate experiments, and representative high-quality images were selected.

### *In vitro* splenocyte stimulation assays

Spleens from OT-I mice were harvested and processed into single-cell suspensions through a 70-micron cell strainer (Grenier). Red blood cells were lysed using ACK Lysis Buffer (Gibco). Splenocytes were plated in 96 well TC-treated U bottom plates (Falcon) at a concentration of 300,000 cells/well in RPMI supplemented with 10% FBS (Gibco), 100 U/mL Penicillin-Streptomycin (Gibco), 1x sodium pyruvate (Thermo Fisher Scientific), 1x nonessential amino acids (Thermo Fisher Scientific) and 1x 2-mercaptoethanol (Thermo Fisher Scientific). Cells were pulsed with αCD45 for 30 minutes at 37°C, washed twice in complete media, and resuspended in 200 μL XL or polyclonal proteins at the indicated concentrations. OVA peptide (SIINFEKL, Genscript) or G4 peptide (SIIGFEKL, Genscript) was added to wells at the indicated final concentrations. Splenocytes were cultured for 48 hours, before harvesting for downstream flow cytometry analysis.

### Primary CD8^+^ T cell isolation

Spleens from C57BL/6J mice were harvested and processed into single-cell suspensions through a 70-micron cell strainer (Grenier). CD8^+^ T cells were isolated using the EasySep^TM^ Mouse CD8^+^ T cell isolation kit (StemCell Technologies) and resuspended at a concentration of 10^6^ cells/mL in RPMI supplemented with 10% FBS (Gibco), 100 U/mL Penicillin-Streptomycin (Gibco), 1x sodium pyruvate (Thermo Fisher Scientific), 1x nonessential amino acids (Thermo Fisher Scientific) and 1x 2-mercaptoethanol (Thermo Fisher Scientific). Medium was additionally supplemented with 10 ng/mL of murine IL-2 (BioLegend) before resuspension and subsequent passaging. Isolated CD8^+^ T cells were activated for 48 hrs. on a non-tissue culture-treated plate that was precoated with 1 μg/mL of αCD3 (BioXCell, clone 2C11) and 10 μg/mL of αCD28 (BioXCell, clone 37.51) in PBS overnight at 4 °C. After activation, T cells were cultured for 48 hrs. in complete media supplemented with murine IL-2 before use in downstream experiments.

### Phosphorylation analysis by flow cytometry

Primary CD8^+^ T cells cultured as described above were starved of IL-2 for 24 hrs. and seeded into 96-well plates at 100k cells/well. Cells were pulsed with αCD45 for 30 minutes at 37°C, washed twice in complete media, and resuspended in 4°C media. Cells were stained with XL on ice for 15 min. followed by 3 min. at 37°C. Cells were immediately fixed with an equal volume of BD Phosflow Fixation Buffer I (BD Biosciences) at 37°C for 10 minutes. Cells were permeabilized for 30 minutes on ice with BD Phosflow Perm Buffer III (BD Biosciences) that had been pre-chilled to-20C. Subsequent washes and surface staining was performed in PBS supplemented with 1% bovine serum albumin (Sigma) and 2mM EDTA (Invitrogen). Samples were resuspended in Fc block (clone 93, eBioscience, 1:50) prior to surface staining for 15 minutes at 4°C. Antibodies against surface targets were stained at 4°C overnight. Cells were analyzed using a BD FACSymphony A3 and data was analyzed in FlowJo v10.

### Förster resonance energy transfer assays

Recombinant αCD45 was conjugated with donor fluorophore Alexa Fluor 555, and a separate αCD45 aliquot conjugated with acceptor fluorophore Alexa Fluor 647, using N-hydroxysuccinimide (NHS) ester chemistry (Invitrogen). Proteins were mixed and dye-matched to a 1:1 molar ratio of donor: acceptor fluorophore. RAW264.7 cells or murine splenocytes were pulsed with 100 nM donor, acceptor, or mixed donor/acceptor αCD45 for 30 min at 37°C in 96 well plates. Cells were washed twice with complete DMEM and treated with titrations of XL for 30 min at 37°C. FRET signal was measured in the PE-Cy5 channel (561 nM excitation laser, 670/30 nM emission filter) using a BD FACSymphony A3, and data was analyzed in FlowJo v10.

Fixed cells were imaged on a Leica Sp8 confocal with a 63x oil objective. FRET efficiencies (E) after crosslinker addition were measured using the FRET acceptor photobleaching method. Briefly, the cells were imaged in the donor and acceptor channels, followed by 90 seconds of acceptor photobleaching (100% laser power) of a selected region of interest. After acceptor bleaching, the cells were reimaged in both donor and acceptor channels. Single color controls and non-crosslinking controls were included to evaluate channel bleed through. FRET efficiency was calculated based on the change in donor emission with the following equation:

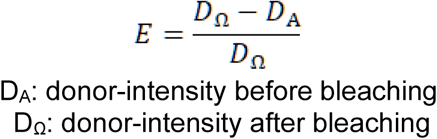

D_A_: donor-intensity before bleaching D_Ω_: donor-intensity after bleaching

### Internalization assays

Recombinant αCD45 was conjugated with Alexa Fluor 647 using N-hydroxysuccinimide (NHS) ester chemistry (Invitrogen). Primary CD8^+^ T cells were isolated as described above and seeded in a 96-well plate and incubated with αCD45 at 10 μg/mL staggered at desired time points. Wells were then split such that one set was incubated with 20 U/ml of ProteinaseK (New England Biolabs) at 37°C for 20 min, while the other set remained at 4°C. AlexaFluor 647 signal was measured via flow cytometry and internalization assays were analyzed as described previously^95^. Briefly, surface fluorescence was calculated as the difference in MFI between untreated and ProteinaseK treated samples, divided by the quenching efficiency of the *t=0* no-internalization control. GraphPad Prism v10 was used to fit a linear regression of the internal fluorescence intensity versus the integral of the surface fluorescence, using the trapezoidal rule to calculate. The slope of the linear regression fit represents the internalization rate (k_e_) and the surface half-life of internalization was calculated as ln(2)/k_e_.

### Tumor inoculation and treatment

Mice aged 6-9 weeks old were injected subcutaneously with 10^6^ tumor cells (B16F10 or MC38) in 50 μL PBS in the shaved right flank. Prior to treatment, mice were randomized to ensure equal mean initial tumor area across groups. αCD45 (40 μg) and XL (60 μg) were treated intratumorally in 20 μL PBS, unless otherwise noted. For alum IL-12 treatment, 5 μg IL-12-ABP and 100 μg alum were mixed in a total volume of 30 μL TBS, incubated at RT for 30 min, and dosed intratumorally on day 6, unless otherwise noted. Tumor area was calculated as the product of tumor length and width (mm). Mice were euthanized when tumor area exceeded 100 mm^2^, or when significant tumor ulceration was observed and recommended by an attending veterinarian.

### Depletion studies

Immune cell depletions were carried out in C57BL/6 mice with antibodies targeting CD8β (BioXCell, Clone 53-5.8, 400 μg twice weekly), NK1.1 (BioXCell, clone PK136, 400 μg twice weekly), Ly6G (BioXCell, clone 1A8, 400 μg twice weekly), and CSF1R (BioXCell, clone AFS98, 300 μg every other day) as previously described^97^. All depletions were dosed i.p. in 100 μL of PBS. Depletions were initiated 1 day prior to treatment and carried out for four weeks.

### Tissue processing for flow cytometry

B16F10 tumors were harvested, weighed, and subsequently minced using dissection scissors in 2.5 mL RPMI with 100 mg/mL Collagenase D (Sigma) and 20 mg/mL DNase I (Sigma). Minced tumors were processed on a gentleMACS Octo-dissociator with heaters (Miltenyi) using program mTDK_1 for B16F10. Dissociated tumors were then filtered through a 70-micron strainer and 25 mg tumor was plated for downstream staining. Tumor-draining inguinal lymph nodes (tdLN) were harvested, weighed, manually dissociated, and filtered through a 5 mL round-bottom tube with cell-strainer cap (Falcon) using the blunt rubber end of a 1mL syringe plunger (BD Falcon). 5 mg of LN was plated for downstream staining. Spleens were harvested, weighed and manually dissociated through a 70-micron strainer using the end of a 1mL syringe plunger (BD Falcon) and red blood cells were lysed using ACK Lysis Buffer (Gibco). Blood was collected via intracardiac puncture in MiniCollect K2-EDTA tubes (Greiner) and red blood cells were lysed using ACK Lysis Buffer (Gibco). Precision counting beads (50uL, BioLegend) were added after initial resuspension and used for downstream data analysis. Viability was assessed with Zombie UV dye (BioLegend, 1:1000) in PBS for 20 minutes at room temperature. Subsequent washes and surface staining was performed in PBS supplemented with 1% bovine serum albumin (Sigma) and 2mM EDTA (Invitrogen). Samples were resuspended in Fc block (clone 93, eBioscience, 1:50) prior to surface staining for 15 minutes. Antibodies against surface targets were stained on ice for 30 min and specific clones are listed in Supplementary Tables 2-6. Cells were fixed with BD Cytofix (BD Biosciences) for 30 min at 4°C. When intracellular staining was performed, cells were fixed and permeabilized with eBiosciences FoxP3 / Transcription Factor Staining Set (ThermoFisher). Cells were assessed using a BD FACSymphony^TM^ A3 and data analysis was performed in FlowJo v10.

### Biodistribution serum measurements

Proteins were labeled with AlexaFluor 647 (ThermoFisher) using NHS-ester conjugation chemistry, as per the manufacturer’s instructions. Blood was collected via intracardiac puncture and serum was isolated using centrifugation through Serum Gel tubes (Sarstedt). Serum fluorescence was measured in black flat bottom 384 well plates (Grenier) on an Infinite M200 microplate reader (Tecan). Free protein concentrations were calculated based on a standard curve prepared using known amounts of serially diluted labeled proteins.

### IFN-γ ELISPOT

ELISPOT plates (BD Biosciences) were pre-coated with IFN-γ capture antibody (BD Biosciences). C57BL/6 mice (n = 5 animals per group) were inoculated with 10^6^ B16F10 cells and treated according to the treatment regimen in **Fig. 3c**. On day 14, 2 days after the end of treatment), spleens were harvested, weighed and manually dissociated through a 70-micron strainer using the end of a 1mL syringe plunger (BD Falcon) and red blood cells were lysed using ACK Lysis Buffer (Gibco). Cells were resuspended in RPMI supplemented with 10% FBS, 1X sodium pyruvate (Gibco), 1X nonessential amino acids (Gibco), 1X beta-mercaptoethanol (Gibco), 100 U/mL Penicillin-Streptomycin (Gibco). B16F10 cells treated with 500 U/mL IFN-γ (Peprotech) overnight were subjected to 120 Gy radiation and dissociated with CellStripper (Corning) into a single-cell suspension in the same complete RPMI. 25,000 irradiated B16F10 cells were co-cultured with 250,000 splenocytes per sample and seeded in a 96-well ELISPOT plate. Plates were incubated for 24 hrs. at 37°C. 5% CO_2_, then developed according to the manufacturer’s protocol. Plates were scanned using a CTL-ImmunoSpot Plate Reader, and data were analyzed using CTL ImmunoSpot Software.

### Tissue processing for scRNAseq

Tumor-draining inguinal lymph nodes (tdLN) were harvested, manually dissociated, and filtered through a 5 mL round-bottom tube with cell-strainer cap (Falcon) using the blunt rubber end of a 1mL syringe plunger (BD Falcon). All lymph node material was used in FACS sorting. Viability was assessed with Zombie Aqua dye (BioLegend, 1:1000) in PBS for 20 minutes at room temperature. Subsequent washes and surface staining were performed in PBS supplemented with 1% bovine serum albumin (Sigma). Samples were resuspended in buffer containing Fc block (clone 93, eBioscience, 1:50) and AlexaFlour 647 labeled p15E tetramer (H2-Kb MuLV env 574-581, NIH Tetramer Core) at 1:100 for 30 min at RT. Extracellular staining was performed at 4°C for 30 min. Cells were also stained with Totalseq C anti-mouse hashing antibodies (BioLegend, 1:100) before FACS. For p15E^+^ CD8+ T cell sorting cells were sorted on a Sony MA900 and gated on FSC and SSC, Live, CD45^+^, CD8^+^, CD44^+^, and p15E Tetramer^+^. Cells were subsequently processed according to the 10x Genomics 5‘ Immune Profiling v3 protocol.

### scRNAseq analysis

The gene expression libraries, VDJ libraries, and cell hashing libraries were pooled according to the manufacturer’s recommendation and sequenced by Illumina NovaSeq 6000 SP. The gene expression and VDJ libraries were aligned to the GRCm39 reference genome and quantified (including TCR clonality) using the Cell Ranger Multi pipeline v9.0.1. The cell hashing sequence reads were processed and quantified using Cell Ranger Count v9.0.1. The gene expression count matrix, TCR contig files, and cell hashtag oligo (HTO) count matrix were processed and analyzed using R v4.3.0, Seurat v5.0.1, and ggplot2 v3.5.1.

The gene expression count matrix was processed using the Seurat (v5.0.1) package in R. The initial quality control filtered out genes that were detected in fewer than 3 cells and removed cells with fewer than 500 genes or greater than 15% mitochondrial genes. Cells were normalized using the NormalizeData() function. The HTO count matrix was added to the Seurat object and normalized. The HTODemux() function was used to assign HTO to each cell. Only singlet cells by HTO assignment were kept for downstream analysis. The cell cycle was predicted using the CellCycleScoring() function, and variable genes were identified using the FindVariableFeatures() function. The ScaleData() function was used to regress out RNA feature counts and percent of mitochondrial genes before performing principal component analysis (PCA) using the RunPCA() function. Batch correction was performed using the CCA integration method within the IntegrateLayers() function. For the initial cell lineage analysis, thirty principal components (PCs) and 500 decision trees were used for constructing the nearest-neighbor graph with the FindNeighbors() function. Thirty neighboring points and twenty PCs were used to generate uniform manifold approximation and projection (UMAP) with the RunUMAP() function. Unsupervised clustering was determined using Louvain clustering as implemented in the FindClusters() function. The clusters with high expressions of Cd3, Trbc, Trac, Cd8, or Cd4 were subset for further analysis. The unsupervised clustering was repeated, clusters with low cell quality (low RNA feature count and high percent of mitochondrial genes) or strong signs of markers from different cell lineages (e.g., Cd3+ Cd19+). The final quality-controlled T cells were clustered using fifty principal components (PCs) and 500 decision trees; the UMAP was generated using fifty PCs, forty neighboring points, and 0.4 minimum distance. Differential gene expression analysis was performed using FindMarkers() and FindAllMarkers() functions with the Wilcoxon Rank Sum test, and the p-values were adjusted using Bonferroni correction.

### BLISS drug independence analysis

Statistical models for estimating drug synergy were developed using the established Bliss independence framework and adapted for Kaplan-Meier survival analysis^55^. Briefly, when treatments A and B are administered individually, they produce survival curves S_A_(t) and S_B_(t), respectively. According to the Bliss independence model, if the drugs act independently, the expected combined survival curve is:

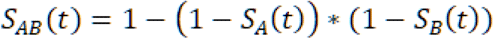

If drugs/treatments are synergistic, the observed drug combination survival will exceed *S_AB_*(*t*). To evaluate drug synergy, we compared the experimental combination survival curve with the calculated drug independence *S_AB_*(*t*) survival curve using the log-rank Mantel-Cox test.

## Statistical analysis

Statistics were performed with GraphPad Prism software V10. As described in figure captions, other metrics were compared by one-way or two-way analysis of variance (ANOVA) with Tukey’s multiple comparisons test, unpaired *t* test, Fisher’s exact test for significant proportions, or the two-sided Wilcoxon rank sum test.The *n* and *p* values are indicated in captions.

## Acknowledgements

We thank the Division for Comparative Medicine (DCM) at MIT for support in animal husbandry and care during animal studies. We also thank the Koch Institute Swanson Biotechnology Center (National Cancer Institute Grant P30-CA14051) for technical support, specifically the flow cytometry, microscopy and histology core facilities, as well as the BioMicro Center. This work was in part supported by National Institute of Biomedical Imaging and Bioengineering (NIBIB) grant no. EB031082, an Aspire Award from the Mark Foundation for Cancer Research and the Koch Institute Support (core) grant no. P30-CA14051 from the National Cancer Institute. L.D. is supported by a graduate fellowship from the Ludwig Center at MIT’s Koch Institute. W.P.III, O.T.P. and J.R.P. are supported by the National Science Foundation (NSF) Graduate Research Fellowship Program. D.J.I. is an investigator of the Howard Hughes Medical Institute.

## Author Contributions

L.D. and K.D.W. conceived, designed and directed this study. Y.J.Z. conducted RNA-seq analysis. A.R. conducted FRET microscopy and analysis. L.D., W.P.III, E.F., Q.H.Z, O.T.P., L.S., J.A.S and J.R.P performed experiments. L.D., Y.J.Z. and A.R. performed data analysis. L.D. and K.D.W. wrote the manuscript, with input from all authors.

