## Extended Data Figures for "CD45 sequestration lowers the signaling threshold in lymphocytes and enhances anti-tumor immunity"

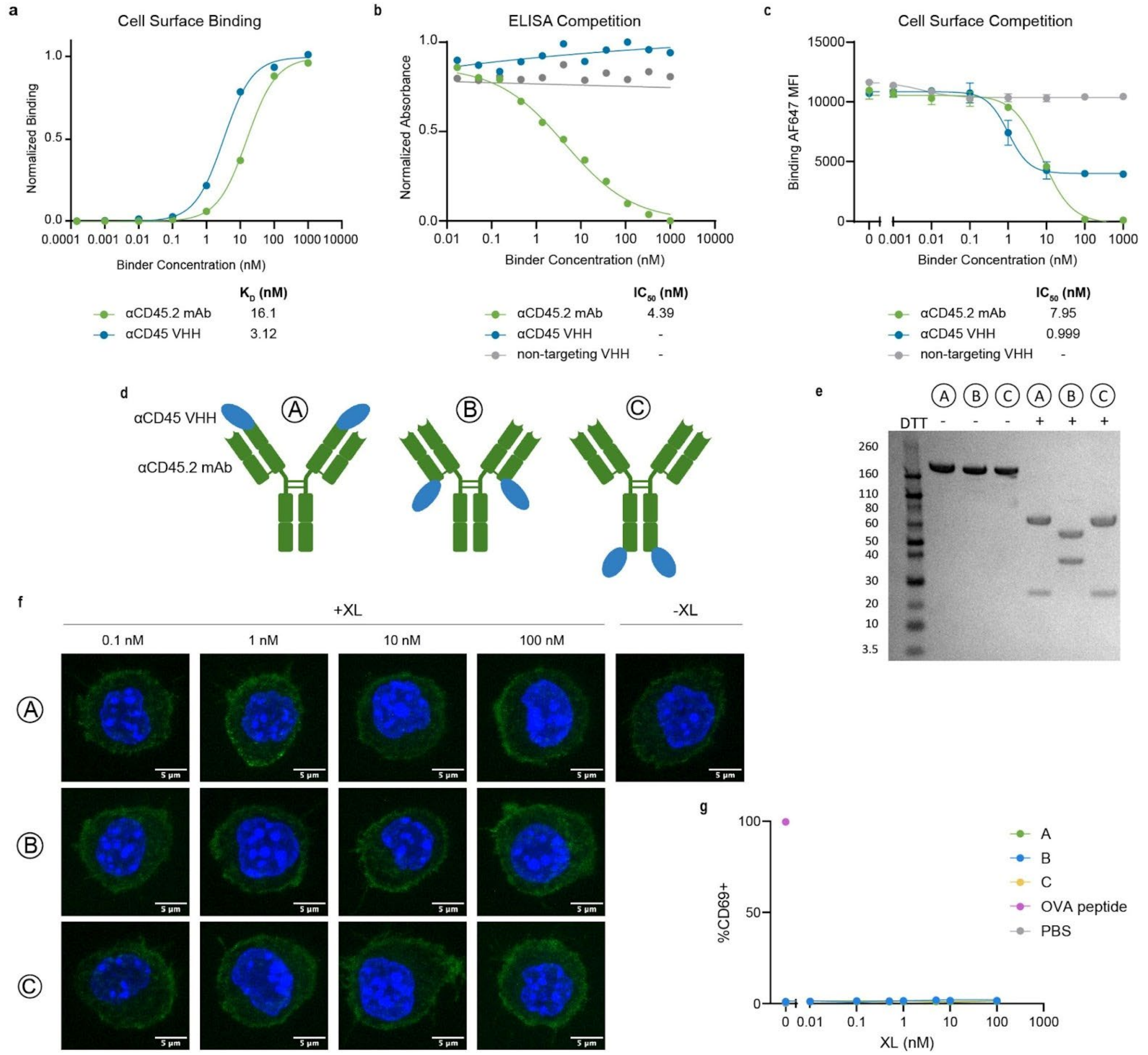


**Extended Data Figure 1. Biepitopic CD45 binding constructs do not crosslink CD45 a.** Equilibrium binding curves normalized to $B_{max}$ of AF647 labeled aCD45 VHH or αCD45.2 IgG on murine splenocytes (n=3 ± s.d.). Equilibrium dissociation constants ($K_D$) were calculated using a nonlinear regression fit for one-site total binding with no nonspecificity. **b.** Competitive ELISA detecting epitope competition between αCD45.2 IgG and αCD45 VHH clones. Biotinylated αCD45.2 IgG binding was detected by streptavidin-HRP, and normalized by maximum absorbance. **c.** Competitive surface binding on murine splenocytes detecting epitope competition between αCD45.2 IgG and αCD45 VHH clones. AF647 labeled αCD45.2 IgG binding was detected by flow cytometry. Inhibitory dose-response curves and the half-maximal inhibitory concentrations ($IC_{50}$ values) were calculated assuming a one-site, reversible competition model (mean ± SD; n = 3). **d.** Schematic of αCD45 VHH – αCD45.2 IgG fusions. **e.** αCD45 VHH – αCD45.2 IgG fusions were cloned and produced in the HEK293F expression system. ProteinA purified fractions are shown on SDS PAGE. **f.** Confocal Z-stack projections of RAW264.7 CD45-mGreenLantern treated with titrations of αCD45 VHH – αCD45.2 IgG fusions for 30 min. CD45 is shown in green and nuclei in blue. **e.** OT-I T cell activation was assessed as shown in Fig. 2e. Percent of CD69+ OT-I T cells, measured at various concentrations of the indicated αCD45 VHH – αCD45.2 IgG fusions.

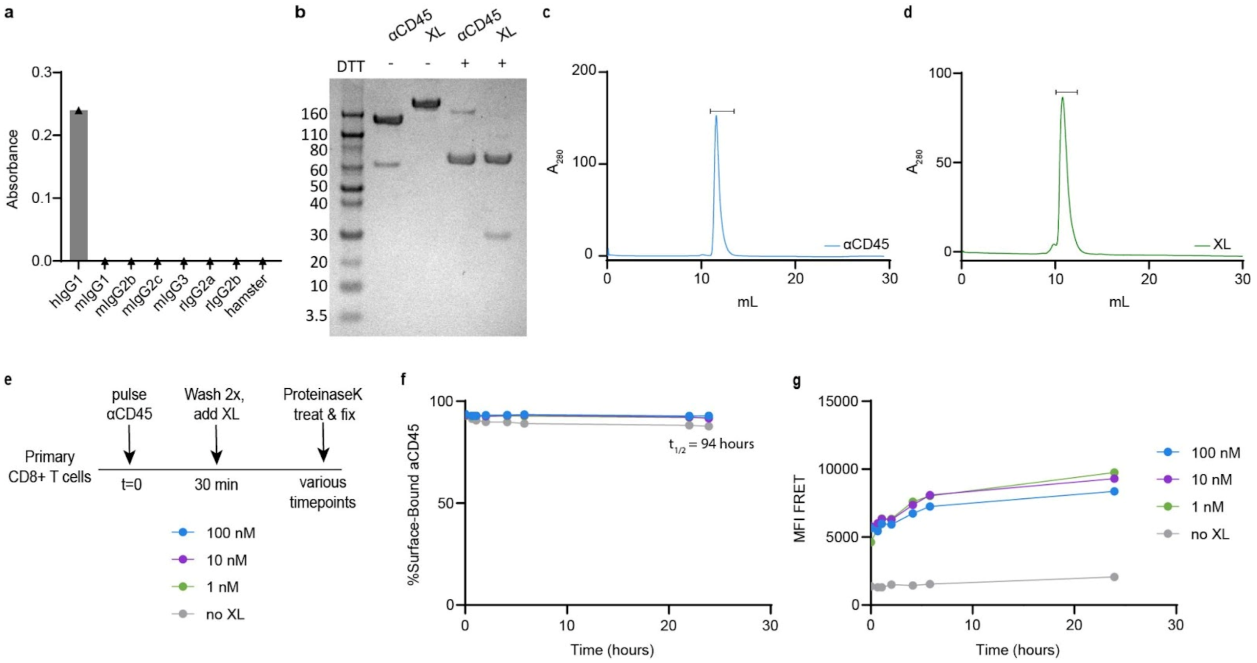


**Extended Data Figure 2. Engineered multivalent binding constructs can cluster CD45 without driving internalization. a.** Equilibrium binding ELISA detecting αhIgG VHH reactivity to various immunoglobulin isotypes. Biotinylated αhIgG VHH binding was detected with streptavidin-HRP at 450 nm absorbance. **b.** αCD45 and XL constructs were cloned and produced in the HEK293F expression system. ProteinA and Nickel-NTA purified fractions are shown on SDS PAGE. **c-d.** Representative size exclusion chromatography and purified fractions for αCD45 (**c**) and XL (**d**) constructs. **e.** Experimental design for aCD45 internalization assay and FRET time-course in **f-g. f.** Internalization kinetics of AF647 labeled αCD45 in the presence or absence of XL after binding to primary activated $CD8^+$ T cells. Surface signal was calculated by ProteinaseK treatment. **g.** FRET kinetics of mixed AF555 and AF647 labeled αCD45 in the presence or absence of XL after binding to primary activated $CD8^+$ T cells. FRET signal was measured in the PE-Cy5 channel (561 nM excitation laser, 670/30 nM emission filter) via flow cytometry.

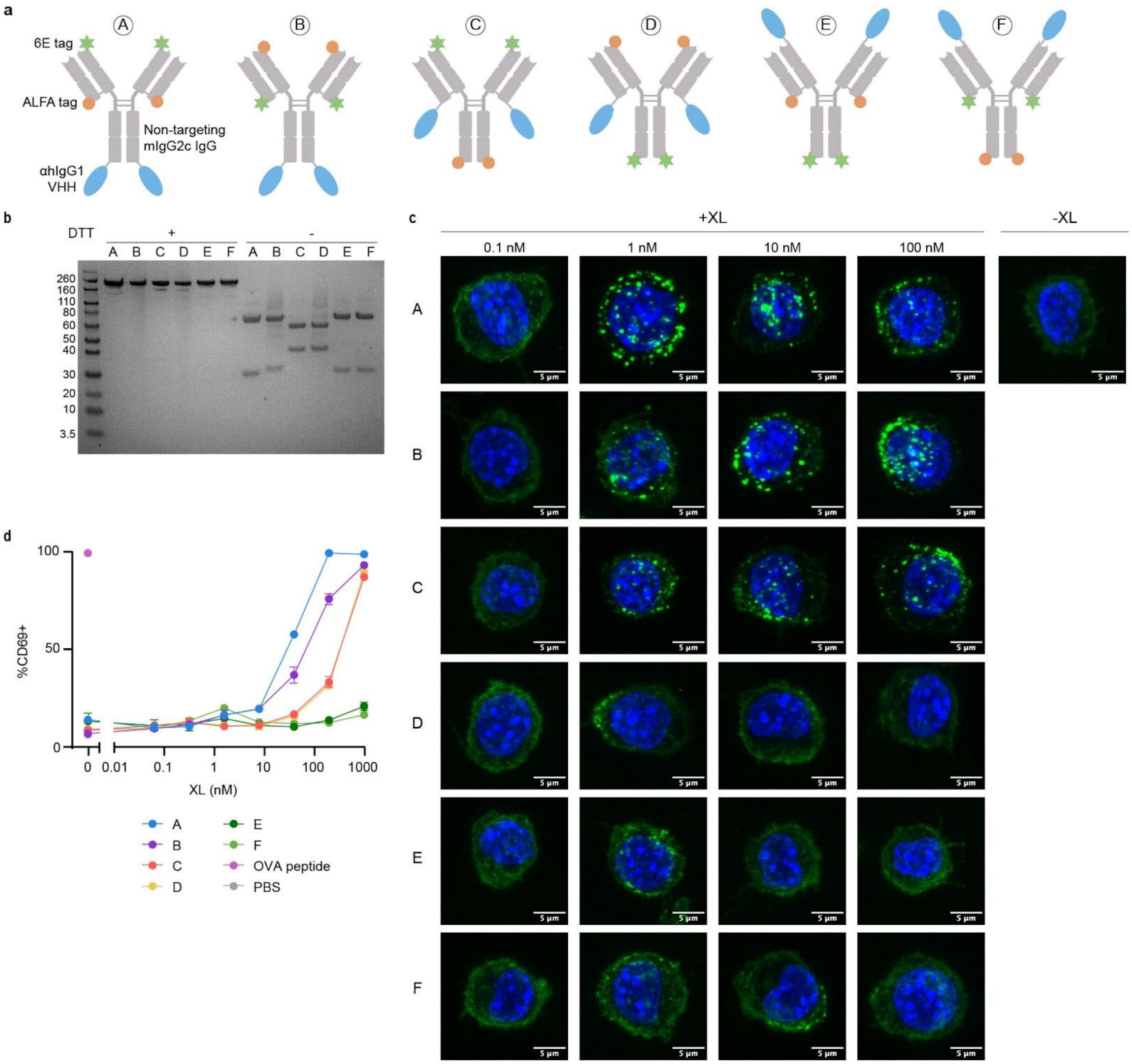


**Extended Data Figure 3. Orientation of crosslinking domains influences CD45 cluster formation a.** Schematic of multivalent XL fusion proteins, constructed by fusing indicated nanobody or peptide tags to an irrelevant anti-FITC IgG as a scaffold. **b.** Multivalent XL constructs were cloned and produced in the HEK293F expression system. Nickel-NTA purified fractions are shown on SDS PAGE. **c.** Confocal Z-stack projections of RAW264.7 CD45-mGreenLantern pulsed with 100 nM αCD45 for 30 min and treated with titrations of XL. CD45 is shown in green and nuclei in blue. **d.** OT-I T cell activation was assessed as shown in Fig. 2e. Percentage of CD69+ OT-I T cells was measured for indicated multivalent XL constructs (mean ± SD; n = 3).

.

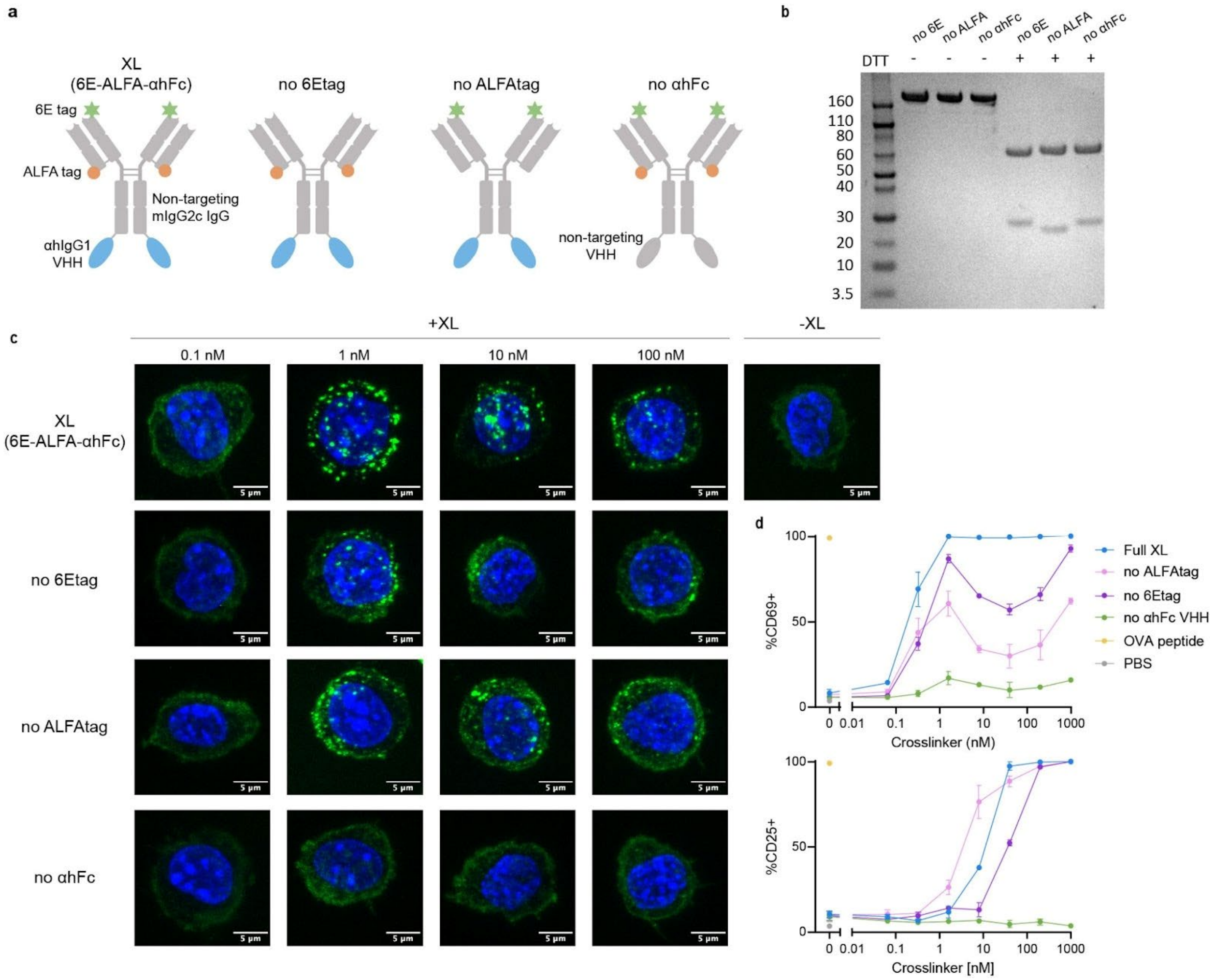


**Extended Data Figure 4. The number of crosslinking domains influences CD45 cluster formation a.** Schematic of multivalent XL fusion proteins, constructed by fusing indicated nanobody or peptide tags to an irrelevant anti-FITC IgG as a scaffold. **b.** Multivalent XL constructs were cloned and produced in the HEK293F expression system. Nickel-NTA purified fractions are shown on SDS PAGE. **c.** Confocal Z-stack projections of RAW264.7 CD45-mGreenLantern pulsed with 100 nM αCD45 for 30 min, and treated with titrations of XL. CD45 is shown in green and nuclei in blue. **d.** OT-I T cell activation was assessed as shown in Fig. 2e. Percentage of CD69+ or CD25+ OT-I T cells was measured for indicated multivalent XL constructs (mean ± SD; n = 3).

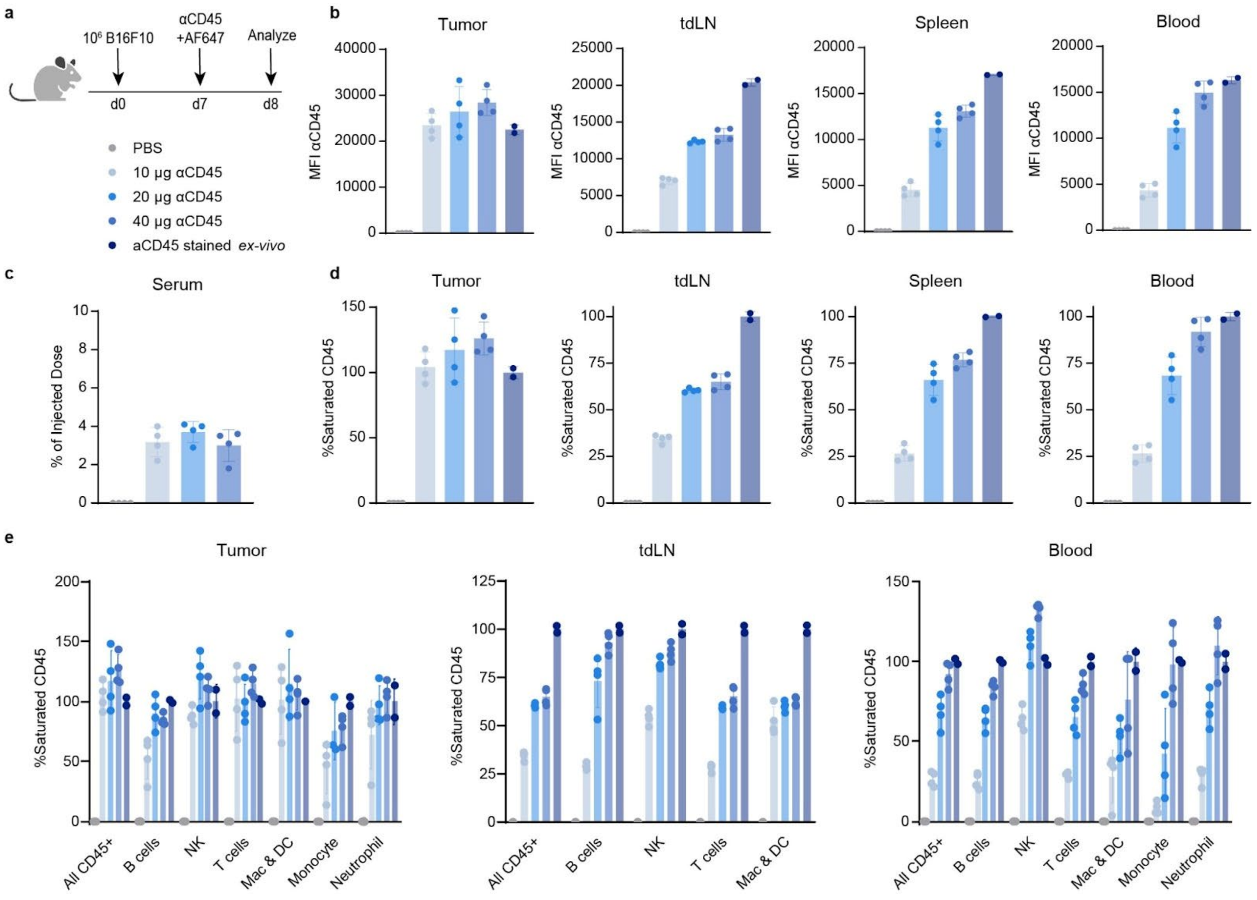

**Extended Data Figure 5. Biodistribution of αCD45 *in vivo.* a.** C57BL/6 mice were inoculated with 1x10$^6$ B16F10 cells. On day 7, mice were treated *i.t.* with the specified doses of AF647-labeled αCD45. 24 hours later, AF647 signal on CD45+ cells was profiled by flow cytometry in various tissues (mean ± SD; n = 4). Additional tissue from untreated mice was stained *ex*-vivo with 100 nM αCD45 during extracellular staining. **b.** Absolute AF647 MFI of CD45+ cells in various tissues. **c.** Serum quantification of % injected dose of αCD45 based on fluorescence spectroscopy measurements. **d.** Relative quantification of CD45 receptor saturation by dosed αCD45. AF647 MFI is normalized to *ex-vivo* stained control (100% occupied CD45) and untreated (0% occupied CD45). **e.** Absolute AF647 MFI of specific lymphocyte cell types in various tissues.

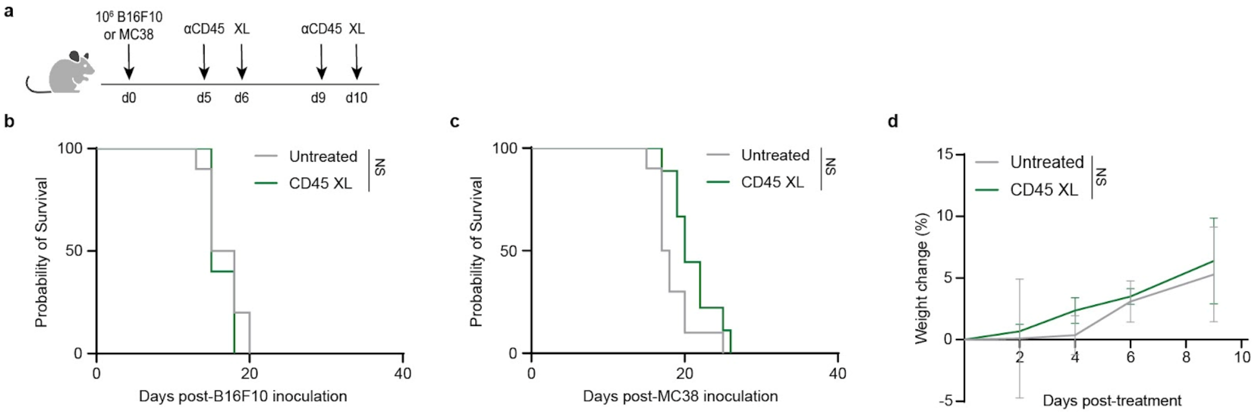


**Extended Data Figure 6. CD45 sequestration is not effective as a monotherapy treatment. a.** Mice (*n* = 5/group for B16F10, n =9-10/group for MC38), inoculated with 1x$10^6$ tumor cells and treated with 40 μg αCD45 and 60 μg XL as shown. **b.** Kaplan Meier survival curve for B16F10 tumors. **c.** Kaplan Meier survival curve for MC38 tumors. **d.** Percent weight change measured from the start of treatment on day 6 (n = 5/group).

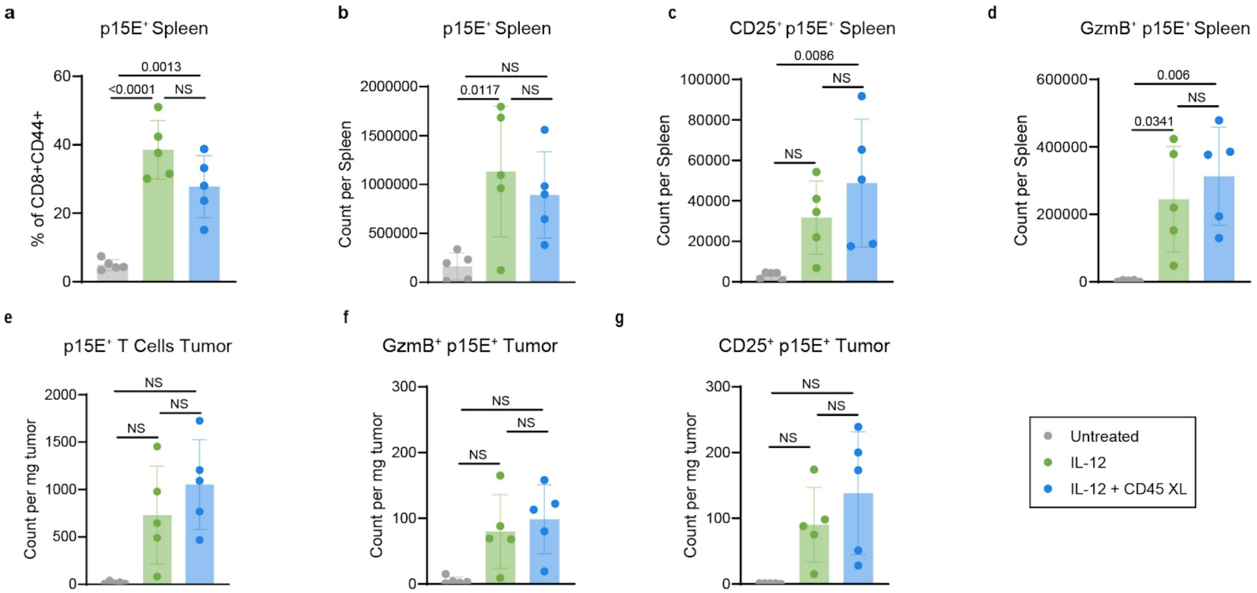


**Extended Data Figure 7. CD45 sequestration combination therapy does not alter tumor-reactive CD8$^{+}$ T cell phenotypes in the spleen or tumor.** C57BL/6 mice (n=5) were inoculated with 1x10$^{6}$ B16F10 cells and treated as shown in Fig. 4a. Tissues were harvested for flow cytometry 48 hours after the completion of treatment. **a.** Proportion of p15E-reactive CD8$^{+}$ CD44$^{+}$ T cells in spleens. **b.** Spleen counts of p15E-reactive CD8$^{+}$ T cells. **c.** Count of CD25$^{+}$ p15E$^{+}$ CD8+ T cells in spleens. **d.** Count of GzmB$^{+}$ Eff/EM CD8+ T cells in spleens. **e.** Tumor counts of CD8$^{+}$ T cells. **f.** Count of CD25$^{+}$ p15E$^{+}$ CD8+ T cells in tumors. **g.** Count of GzmB$^{+}$ p15E$^{+}$ CD8+ T cells in tumors. *p* values were determined by one-way ANOVA followed byTukey's multiple-comparison test.

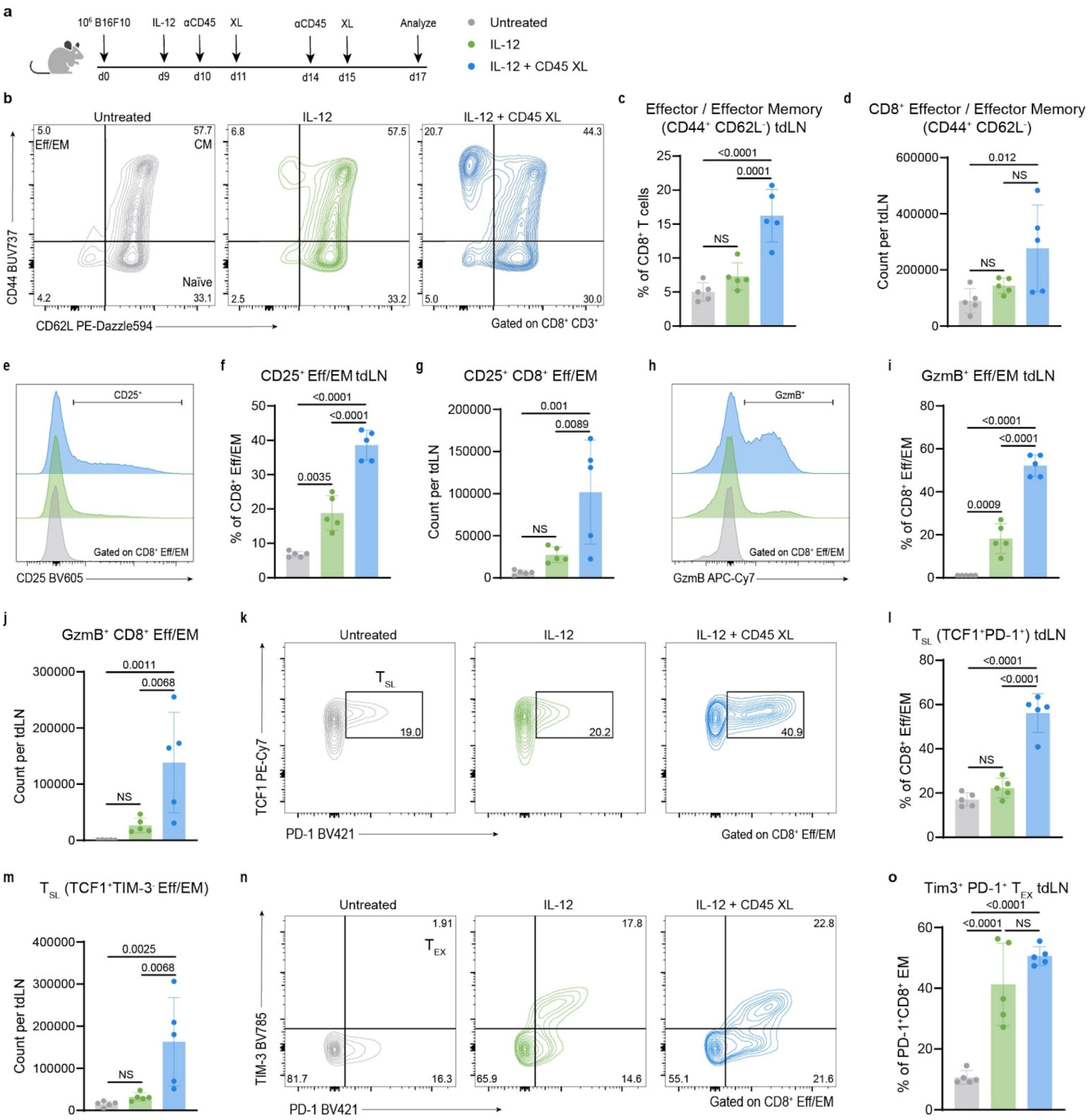


**Extended Data Figure 8. CD45 sequestration combination therapy promotes an activated effector CD8⁺ T cell phenotype in the tdLN. a.** C57BL/6 mice (n=5) were inoculated with 1x10⁶ B16F10 cells and treated as shown. Tissues were harvested for flow cytometry 48 hours after the completion of treatment. **b.** Representative contour plots and gating for effector/effector memory (Eff/EM), central memory (CM), and naïve CD8⁺ T cells in the tdLNs, previously gated on live CD45⁺CD3⁺CD8⁺. **c.** Proportion of Eff/EM CD8⁺ T cells in tdLNs. **d.** tdLN counts of Eff/EM CD8⁺ T cells. **e.** Representative contour plots and gating for CD25 expression in Eff/EM CD8⁺ T cells. **f.** Proportion of CD25⁺ Eff/EM CD8+ T cells in tdLNs. **g.** tdLN counts of CD25⁺ Eff/EM CD8+ T cells. **h.** Representative histogram of granzyme B (GzmB) expression in Eff/EM CD8⁺ T cells. **i.** proportion of GzmB⁺ Eff/EM CD8+ T cells in tdLNs. **j.** tdLN counts of GzmB⁺ Eff/EM CD8⁺ T cells. **k.** Representative stem-like T cell (TCF1⁺PD-1⁺) gating. **l.** tdLN frequency of stem-like PD-1⁺TCF1⁺ Eff/EM T cells. **m.** tdLN counts of stem-like PD-1⁺TCF1⁺ Eff/EM T cells. **n.** Representative exhausted T cell

(PD-1$^+$TIM3$^+$) gating. **o.** tdLN frequency of exhausted PD-1$^+$TIM3$^+$ Eff/EM T cells. *p* values were determined by one-way ANOVA followed by Tukey's multiple-comparison test.

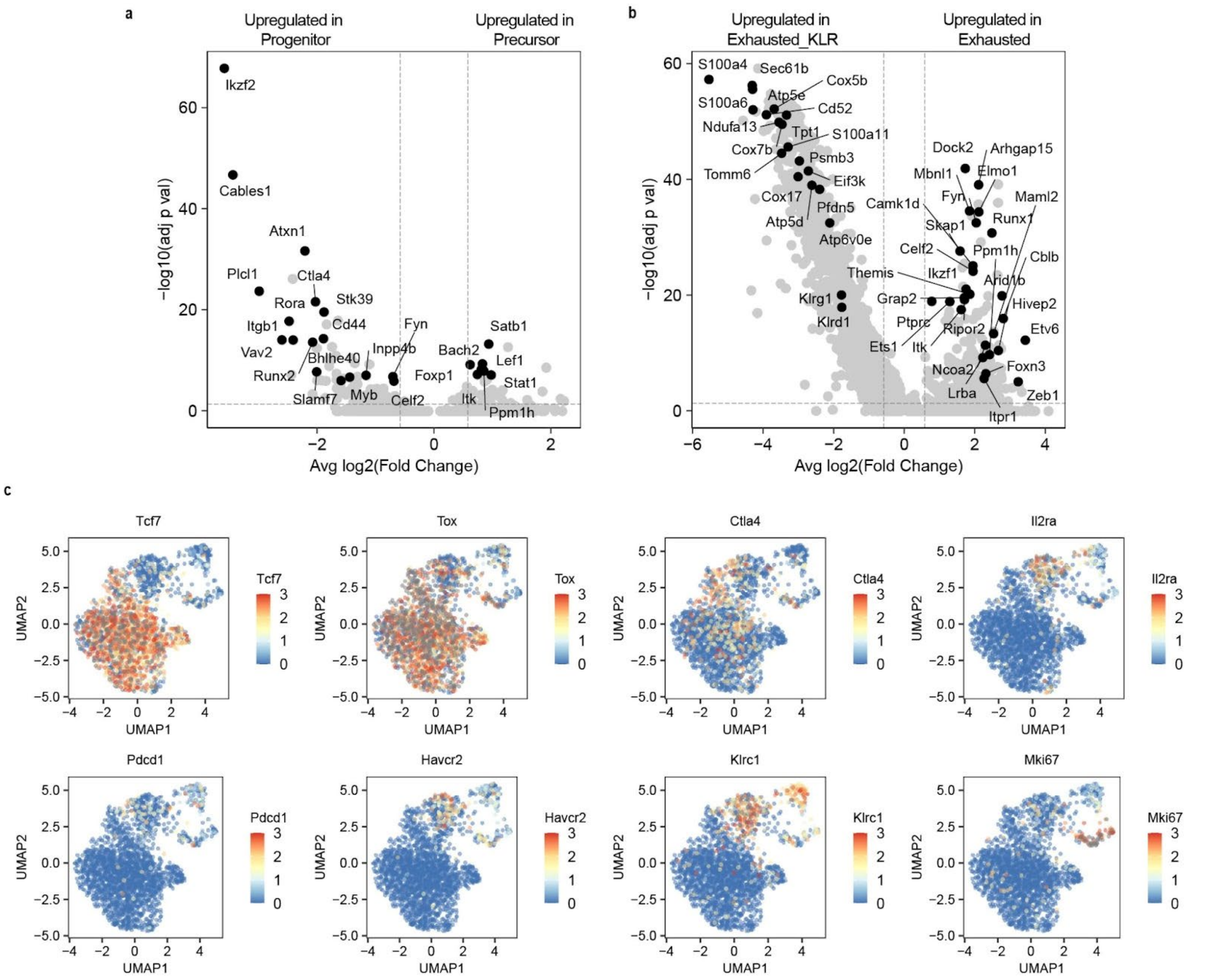


**Extended Data Figure 9. Transcriptomic signatures of phenotypic T cell states. a.** Volcano plot of transcripts differentially expressed between precursor-exhausted T cells and progenitor-exhausted T cells. **b.** Volcano plot of transcripts differentially expressed between terminally exhausted T cells and exhausted KLR T cells. *p* values for volcano plots were calculated using a two-sided Wilcoxon rank sum test and were adjusted using Bonferroni correction. Genes with an average log-fold change >0.25 and an adjusted *p* value $<1\times10^{-5}$ were considered significant. **c.** UMAP of gene expression signatures for selected genes.

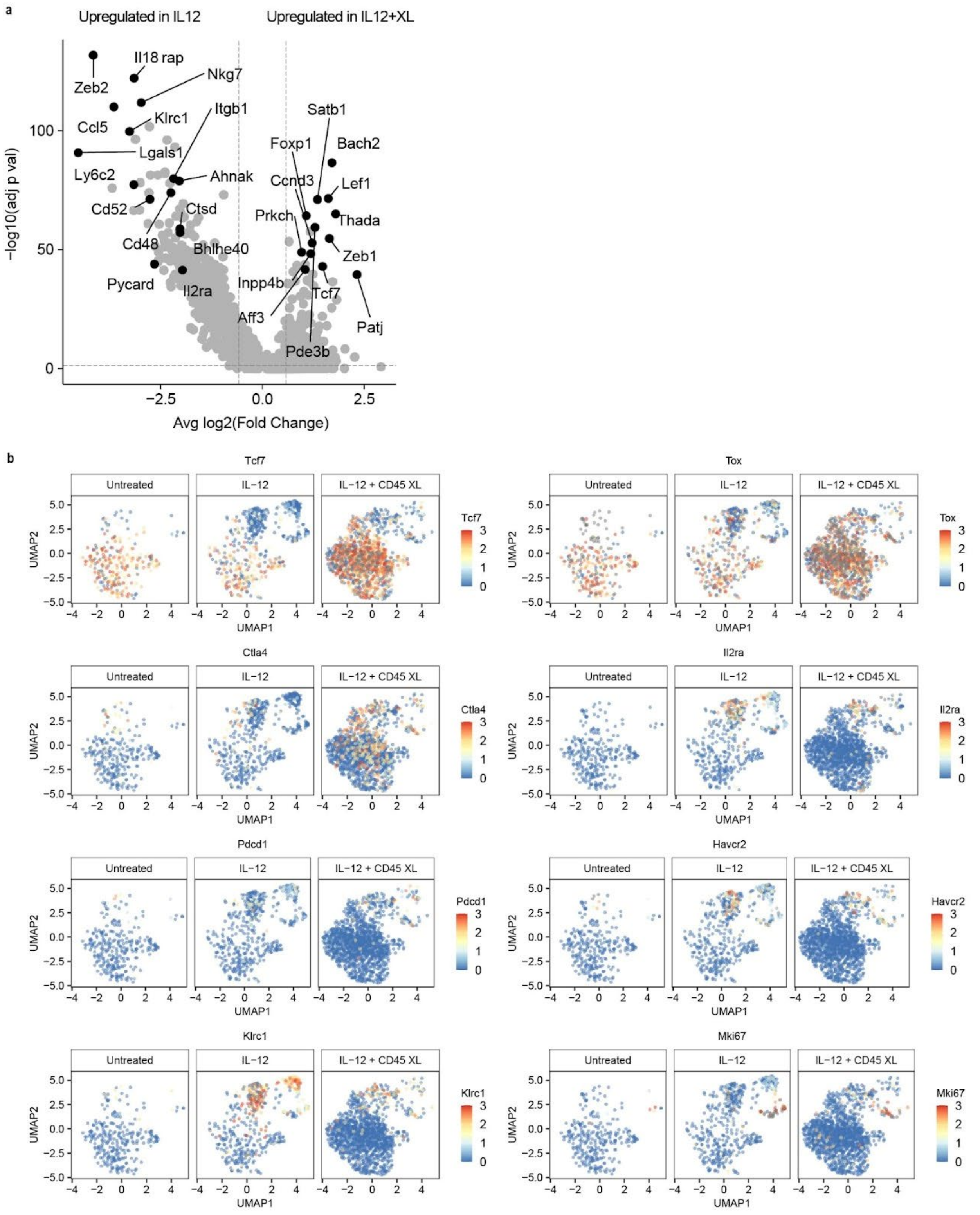


**Extended Data Figure 10. Combination therapy upregulates stemness signatures. a.** Volcano plot of transcripts differentially expressed between p15E-reactive T cells recovered from IL12 and IL12 + CD45 XL groups. *p* values for volcano plots were calculated using a two-sided Wilcoxon rank sum test and were adjusted using Bonferroni correction. Genes with an average log-fold change >0.25 and an adjusted *p* value

$<1\times10^{-5}$ were considered significant. **b.** UMAP of gene expression signatures for selected genes, across treatment groups.
